# Interspecific transfer of genetic information through polyploid bridges

**DOI:** 10.1101/2023.11.20.567804

**Authors:** Felipe Kauai, Quinten Bafort, Frederik Mortier, Marc Van Montagu, Dries Bonte, Yves Van de Peer

## Abstract

Many organisms have more than two sets of chromosomes, due to whole genome duplication (WGD), and are thus polyploid. Despite usually being an ephemeral state in the history of life, polyploidy is widely recognized as an important source of genetic novelty over macroevolutionary scales. More recently, polyploidy has also been shown to facilitate interspecific gene flow, circumventing reproductive barriers between their diploid ancestors. Yet, the implications of WGD-linked introgression on community-level evolutionary dynamics remain unknown. Here, we develop a model of cytotype dynamics within mixed-ploidy populations to demonstrate that polyploidy can in fact serve as a bridge for gene flow between diploid lineages, where introgression is fully or partially hampered by the species barrier. Polyploid bridges emerge in the presence of triploid organisms, which despite critically low levels of viability, can still allow the transfer of alleles between diploid states of independently evolving mixed-ploidy species. Notably, while marked genetic divergence prevents WGD-mediated interspecific gene flow, we show that increased recombination rates can offset these evolutionary constraints, which allows a more efficient sorting of alleles at higher-ploidy levels before introgression into diploid gene pools. Additionally, we derive an analytical approximation for the rate of gene flow at the tetraploid level necessary to supersede introgression between diploids with non-zero introgression rates, which is especially relevant for plant species complexes, where interspecific gene flow is ubiquitous. Altogether, our results illustrate the potential impact of polyploid bridges on evolutionary change within and between mixed-ploidy populations.

## INTRODUCTION

Polyploidy, the outcome of whole genome duplication (WGD), is a mutational event that has pervasive consequences for the evolutionary trajectory of organisms (1-6). Ongoing efforts to recover WGDs in the history of life (7, 8) reveal that several of such events seem to cluster around periods of environmental upheaval (9-11), such as the mass extinction event at the Cretaceous-Paleogene (K-Pg) boundary (12, 13), indicating a potential enhanced capacity of polyploids to adapt and respond to stressful conditions (4, 14-17). Notwithstanding the presumed adaptive potential, phylogenetic and theoretical analyses also suggest that polyploid lineages display greater extinction rates than diploids (18-20), or eventually evolve to the more stable diploid state through the process of rediploidization (11). Despite the ambivalence, WGD is widely recognized to have important implications for the generation of genetic novelty over macroevolutionary scales (3, 9, 21-23). Nevertheless, whether polyploid organisms have an appreciable short-term evolutionary impact on the communities in which they emerge remains unclear.

Recently, gene flow from a well-adapted diploid species, over its tetraploid to the tetraploid of another species, or WGD-mediated adaptive introgression, has been suggested as a potential force in promoting adaptation of polyploid species to specific intracellular demands and challenging environments (24-33). For example, gene flow between *Arabidopsis arenosa* and *A. lyrata* was shown to be facilitated at the polyploid level (31), and seems to have played a major role in stabilizing meiosis among the higher ploidy individuals of both species (27). Introgression mediated by WGD might also have conferred a selective advantage to polyploid amphibians in adapting to extreme environments (28), and have been responsible for reticulate evolution in fungi (34, 35). In fact, artificially induced polyploidy is known among plant breeders to facilitate hybridization of otherwise sterile diploid hybrids, allowing the generation of agriculturally useful species (36, 37). In all respects, these observations support the hypothesis that polyploidy may indeed break the species barrier (28). Still, as there is not a direct pathway for gene flow back to diploids, the phenomenon is expected to operate uniquely at the polyploid level (33, 38).

Triploid cytotypes may offer an indirect route to restore gene flow back to the diploid level, despite their notably low overall viability (39). Indeed, it has been shown that triploids have successful outcrossing rates with different ploidy levels (40). For instance, the discovery that the genes coding for the C_4_ photosynthetic pathway in the tetraploid *Neurachne muelleri* and the diploid, tetraploid and hexaploid cytotypes of *Neurachne munroi* are not paralogous, but likely shared through hybridization or lateral gene transfer (30) suggests that the introgression of alleles through WGD may indeed spread among cytotypes and profoundly impact the evolutionary history of the recipient species. Clear signs of admixture at the polyploid level and intraspecific ploidy variation in other grass lineages, such as *Alloteropsis* (41, 42) suggest that lateral gene transfer, mediated by WGD, does occur (43), and that genetic information can introgress into the (more) stable diploid levels. If this is the case, the duplication of the entire genome of an organism may not only promote the generation of genetic novelty but also the distribution of extant genetic material within ecosystems.

This is particularly relevant as introgression events strongly affect phylogenomic reconstructions of species divergence, especially in plants, where WGD-mediated gene flow is widespread (44). Discerning the routes through which genetic material flows within ecosystems – a process deeply intertwined with population structure and dynamics (45) – can thus contribute to our general understanding of evolutionary change across ecosystems. For instance, if WGD significantly facilitates interspecific introgression, then the potential for intraspecific flow of genetic material across ploidy levels motivates questions concerning the extent with which WGD impacts reproductive isolation and, consequently, speciation rates. Since polyploidy incidence in plants is strikingly common (46, 47), as well as in several animal clades (16), a systematic understanding of gene flow and cytotype dynamics within and among mixed-ploidy systems is paramount.

The current understanding of the mechanisms underlying the emergence and dynamics of polyploidy within diploid populations stems from extensive theoretical and experimental research, which continue to provide valuable insights on the frequency and occurrence of WGD in nature (14, 20, 48-53) . Most models assume that polyploids primarily emerge by the union of unreduced (diploid) gametes, which are formed by a rare event resulting from the non-segregation of chromosomes during meiosis (39, 54, 55). Unreduced gametes may fuse within a population of diploid organisms to give rise to a new tetraploid cytotype, which is then expected to face significant challenges to establishment due to negative frequency-dependent selection; a phenomenon known as *minority cytotype exclusion* (56). As such a path to polyploid formation depends on the joint probability of a very rare event (39), it has been hypothesized that triploid intermediates could relieve the frequency-dependent disadvantage of the emerging tetraploid cytotype by functioning as bridges between ploidy levels (39, 40, 49, 50). This observation is supported by a number of experimental efforts, demonstrating that triploids also promote bidirectional interploidy gene flow in many systems, despite critically low levels of fertility relative to diploids and tetraploids (38, 39, 53, 57-63).

To understand gene flow within mixed-ploidy populations, here we generalize a model for cytotype dynamics introduced by Felber (48) to include partially viable triploids and explicit genetic information. We provide a new analytical account of the model, derive the equilibrium cytotype frequencies and gauge the role of triploids within mixed-ploidy populations as far as its interactions with tetraploid and diploid cytotypes are concerned. Next, we extend this model to a broader theoretical framework consisting of two independent mixed-ploidy populations that are partially connected at the tetraploid level, *i*.*e*., displaying WGD-potentiated hybridization, using a stochastic individual-based simulation. This theoretical construction allows us to measure how alleles flow among cytotypes and between two independent mixed-ploidy populations. We then set out to explore the conditions and constraints for WGD-mediated interspecific gene flow between diploid cytotypes of different mixed-ploidy species and quantify how much gene flow at the tetraploid level is necessary to supersede introgression of an allele with adaptive value between diploid species, whose introgression rates may be non-zero, as is the case for many plant species complexes. Altogether, our results provide insights into the extent to which polyploids can contribute to interspecific gene flow and adaptive introgression, despite their usually transient permanence in ecological systems.

## RESULTS

### Model for cytotype dynamics in a single mixed-ploidy population

We assume an initially diploid population consisting of *N* individuals which reproduce sexually in discrete non-overlapping generations. Specifically, we assume that in each generation an effectively infinite pool of gametes is formed, of which a random sample of gametes is obtained to generate the next generation. Gametes can either be haploid or diploid. When two haploid (diploid) gametes fuse, a diploid (tetraploid) individual is produced, whereas the fusion of haploid and diploid gametes produces a triploid individual. Diploid organisms produce diploid gametes (unreduced) with probability *ν* and haploid gametes with probability (1 − *ν*). The emerging triploid cytotypes are assumed to display reduced viability *φ* and produce haploid or diploid gametes, with equal probability, as studied by previous theoretical models (49, 50) (see Discussion). Tetraploids, on the other hand, are allowed to generate only diploid gametes, as there is no known mechanism of viable haploid gamete formation by tetraploid cytotypes (38) (but see (64) for one record of haploid gamete formation by tetraploids in fish).

We define such a population, which consists of different cytotypes that emerge through the formation and random fusion of different gamete types as a mixed-ploidy population. Notice that because only haploid and diploid gametes are allowed in our model (across all ploidy levels), only three cytotypes can emerge by construction, namely, diploids, triploids, and tetraploids. The emergence of new cytotypes in the population is therefore a function of the random fusion of gamete types and is, as such, conditioned on the cytotypes in the population only. It is instructive to consider the cytotype frequency dynamics in the limit *N* ⟶ ∞, as studied by Felber (48), where we can describe the time evolution of cytotype frequencies within such a mixed-ploidy population through a set of non-linear recursive equations. Let *g*_*p*_ be the frequency of the *p*-ploid gamete in the gamete pool produced, where *p* ∈ {1, 2}. Then, after one generation of random mating one can compute the frequencies of each gamete type as follows:

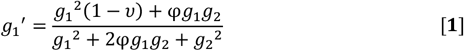

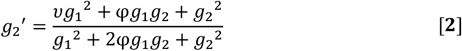

To precisely understand the formulation in Eqs 1 and 2, let X_2_, X_3_ and X_4_ represent the frequency of diploid, triploid and tetraploid cytotypes, respectively. In a mixed-ploidy population which produces haploid and diploid gametes with frequencies g_1_ and g_2_, respectively, the expansion of the binomial (*g*_1_ + *g*_2_)^2^ returns the expected frequency of each cytotype: X_2_ = *g*_1_^2^, X_3_ = 2*g*_1_*g*_2_ and X_4_ = *g*_2_^2^. By reducing the contribution of triploids to the gamete pool by a factor φ, we arrive at the formulation for the dynamics of gametes in Eqs. **1** and **2**. Then, gamete equilibrium frequencies can be obtained by setting *g*_1_^′^ = *g*_1_ and *g*_2_^′^ = *g*_2_, and solving for *g*_1_ (note that *g*_2_ = 1 − *g*_1_). We find that the equilibrium gamete frequencies satisfy:

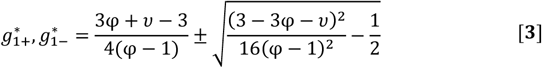

Eq. **3** tells us that two fixed points for *g*_1_ exist for appropriate values of triploid viability φ and unreduced gametes frequency *ν*, one of which is unstable and the other one stable (see *SI Appendix Text S1*). The stable equilibrium frequencies, which the system always converges to under the assumed initial conditions (an initial population of diploids), are shown for each cytotype as a function of *ν* and φ (Fig. 1*A*). The rate at which unreduced gametes are produced by diploids has a direct influence on the frequencies of each cytotype in the population. When φ = 0, i.e., triploids are completely unviable, we observe that tetraploids overtake the system (X_4_ = 1) for *ν* = 0.171. This is the maximum limit of unreduced gametes frequency *ν*^′^ numerically shown by Felber (48) to allow stable coexistence between diploids and tetraploids in the absence of triploids and external selective pressures. Our formulation of Felber’s model allows us to generalize his results to the case where triploids inhabit the system with any viability (φ > 0), and obtain an explicit analytical expression for *ν*^′^ given any φ as follows:

**Fig. 1.**
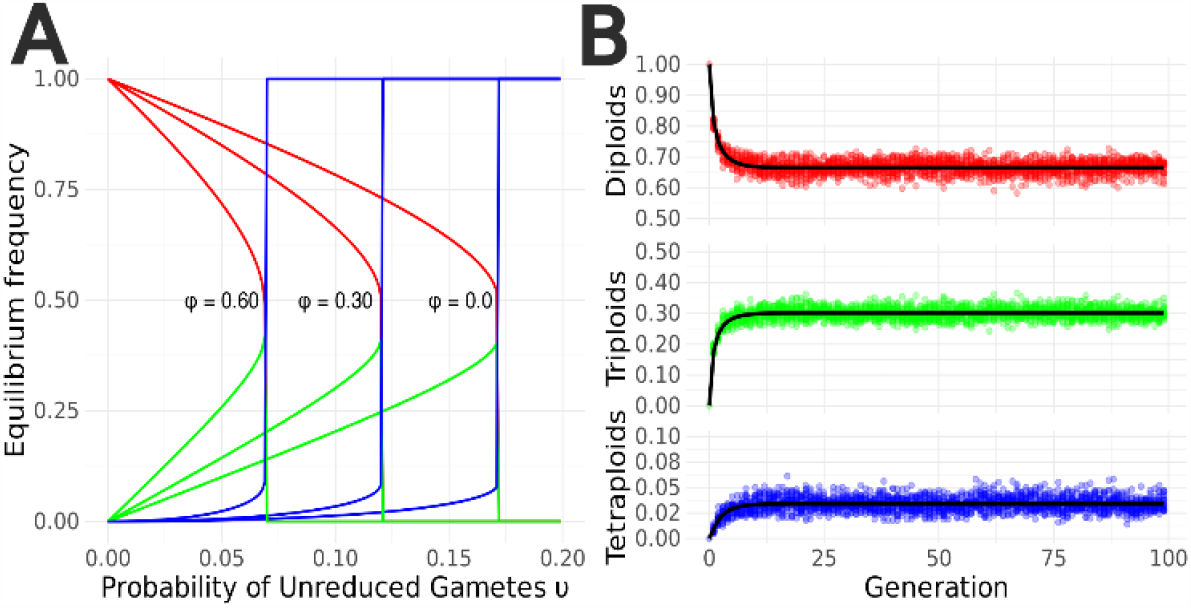
Dynamics of cytotype frequencies in a single deme. (*A*) Equilibrium frequency of diploids (solid red line), triploids (solid green line) and tetraploids (solid blue line) as a function of the probability of unreduced gametes in diploids *ν*, for triploid viability *φ* = 0, *φ* = 0.30 and *φ* = 0.60. Maximum frequency of unreduced gametes 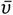 below which coexistence is verified corresponds to the vertical lines, where tetraploids overtake the system. (*B*) Frequency of each cytotype in the mixed-ploidy population per generation, with *φ* = 0.30 and *ν* = 0.10, given by the set of Eqs. **1** *and* **2** in solid black lines, and by the individual-based stochastic version of the model (in faintly colored filled circles) with finite population size *N* = 1000 (see *Methods*).

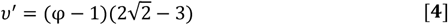

As the probability of unreduced gametes *ν* increases, the frequency of triploid and tetraploid cytotypes becomes progressively higher until a value of *ν*^′^ is reached (see *SI Appendix Text S1* for the proof of the result in Eq. **4**), over which tetraploids dominate. For instance, with triploid viability φ = 0.30, Eq. **4** tells us that *ν*^′^ is 12.01%, and below this threshold cytotypes will coexist, i.e., display stable non-zero frequencies. We confirmed this expectation by numerically iterating Eqs. **1** and **2** and comparing with an individual-based stochastic simulation of the model with finite population sizes (Fig. 1*B*).

The influence of triploid viabilities on the stable frequencies of cytotypes within a population is anticipated by the triploid bridge hypothesis, where triploids may aid the emergence of tetraploids by reducing the effect of *minority cytotype exclusion* (50). To understand it in a qualitative manner, notice the probability that a triploid is formed within a diploid population is 2(1 − *ν*)*ν*, whereas the formation probability of a tetraploid is simply *ν*^2^, with *ν* ≪ 1. As we increase triploid fertility, the frequencies of triploid cytotypes converge to higher magnitudes in the system. By extension, the relative proportion of diploid gametes increases as they are produced with probability *ν* in diploids and probability 0.5 in triploids (Eq. **2**). The higher load of diploid gametes within the population, essentially due to a partial replacement of diploids by triploids, then facilitates the expansion of tetraploid cytotypes. Therefore, despite reduced viability, triploids can relieve the frequency-dependent disadvantage of tetraploids, aiding their establishment by introducing a higher number of diploid gametes in the system (39, 50). This observation is further supported by experimental evidence on the absence of intercytotype post-zygotic barriers in *Hieracium* sp. (53), and direct evidence of intercytotype bidirectional gene flow in *Betula* sp. (60) and *Arabidopsis* sp. (6, 38), for example.

### Mixed-ploidy model of interspecific gene flow

To understand how interspecific gene flow can occur at the diploid level through polyploidization, we have built a stochastic individual-based model of two discrete finite panmictic mixed-ploidy populations. Both populations follow largely independent dynamics, as described in the previous subsection, but are connected by migration with rate *κ* at the tetraploid level, which represents the degree with which tetraploids break the species barrier (28, 33). The initial population consists of a set of identical individuals bearing a diploid genome represented by two finite chromosomes with *L* biallelic unlinked loci. At each mating event, recombination and gamete formation take place (see *Methods*), with offspring emerging from the union of its parents’ gametes. Fig. 2 illustrates the logical structure of the model, whereby two mixed-ploidy species (A and B) are connected only by migration at the tetraploid level. In this work, we analyze interspecific gene flow in a system that has triploids in both mixed-ploidy populations, for conceptual clarity. However, asymmetrical gene flow is also expected, as one of the populations might lack triploid cytotypes due to restrictively low viability. Triploid intermediates are not required for gene flow from diploids to tetraploids but are required for introgression from tetraploids back to diploids (38). A lack of triploid cytotypes in one mixed-ploidy population thus allows for the emergence of incomplete bridges with asymmetrical gene flow only to the diploid in the population with triploids (see *SI Appendix* Fig. S1).

**Fig. 2.**
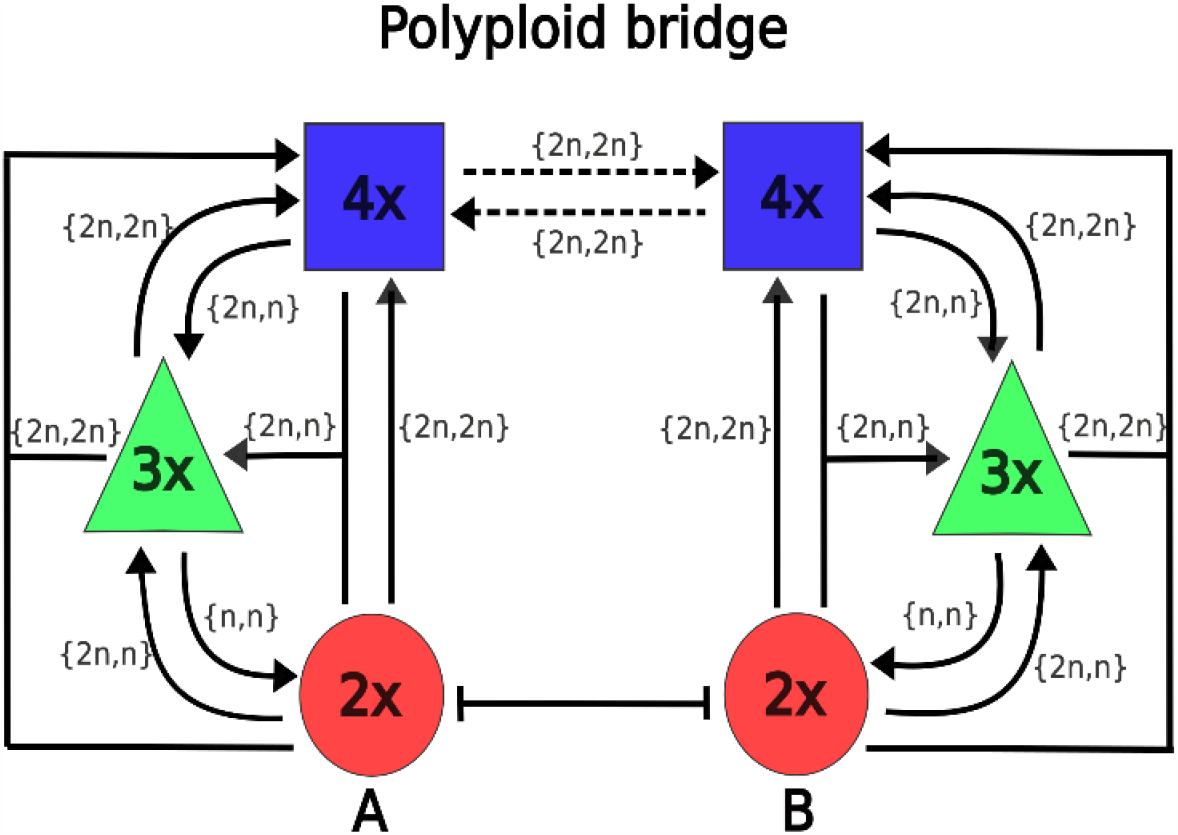
Mixed-ploidy model of interspecific gene flow. The intercytotype gene flow within and between mixed-ploidy populations A and B is depicted, with diploids (2x red circle), triploids (3x green triangle) and tetraploid (4x blue square) cytotypes, following largely independent dynamics, except by migration at the tetraploid level, represented by the dashed lines connecting the tetraploid cytotypes at the upper part of the picture. Solid lines connecting cytotypes represent possible intercytotype mating events, with the direction of the arrow indicating the resulting ploidy level. The unordered set next to each line represents the gametes involved in the mating trial. For example, a 2n (diploid) gamete flowing from a 4x cytotype and a n (haploid) gamete flowing from a 2x cytotype, *i*.*e*., {2n,n}, will result in a 3x cytotype. The solid line between the 2x cytotypes from populations A and B represents the species barrier.

Let *G*_*A*_ and *G*_*B*_ be haploid sequences that represent the reference genome of species *A* and *B*, respectively. Then, we consider a hypothetical divergent lineage, where each mixed-ploidy population represents one of two species that are genetically or ecologically (see Discussion) isolated with genetic distance *d*(*G*_*A*_; *G*_*B*_) (see Fig. 3*A* and *Methods* for technical details). We first assume a neutral environment, i.e., the genotype of each individual has no bearing on how well it performs within its population nor on the genetic compatibility between two reproducing organisms. The process of gene transfer between populations down to diploid levels is then controlled by two parameters: the degree to which polyploids from one population enter the gene pool of the next population, represented by *κ*, and triploid viability φ. The system is initialized so that in each population diploid individuals are fixed for a different allele in each one of the *L* loci (see *SI Appendix Algorithm S1* for technical details on code implementation). Then, as time unfolds, migration at the tetraploid level carries genetic information from one mixed-ploidy population to the next at rate *κ*. We then measured the frequency of introgressed alleles at the diploid state per generation and verified that fixation of non-native alleles can happen even for critically low values of *κ* (Fig. 3*B*).

**Fig. 3.**
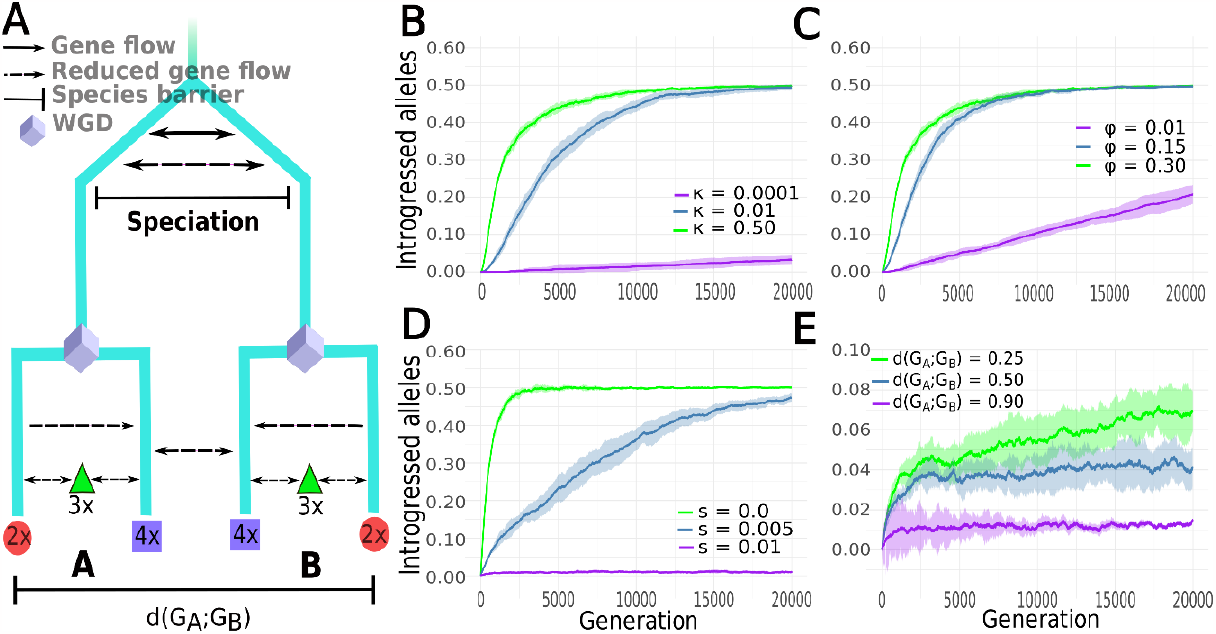
Frequency of introgressed alleles at the diploid level per generation. (*A*) The evolutionary history of a hypothetical lineage splitting into two different species, A and B, is presented with each one displaying WGD events (following (33). Diploids (red circle), triploids (green triangle) and tetraploids (blue square), with the possible gene flow pathways as studied are highlighted. The genetic distance, *d*(*G*_*A*_; *G*_*B*_), represents the nucleotide polymorphisms between the two species following their evolutionary history (see *Methods*). (*B* and *C*) The average frequency of homozygous loci with introgressed alleles per generation is shown, for different values of *κ* and *φ*, respectively, and *φ* = 0.30 (*B*) and *κ* = 0.50 (*C)*. Homozygous and heterozygous loci are distinguished to highlight that fixation proceeds iteratively through successive recombination events, until the system converges to a 50% equilibrium point of introduced and native homozygous loci in each deme (see *SI Appendix* Fig. S5 for the evolution of introgressed alleles considering only heterozygous loci). (D and *E*) The total frequency of introgressed alleles per generation for different selection strength values *s* (with *d*(*G*_*A*_; *G*_*B*_) = 100%) (D) for alternative genetic distances using a selection strength of *s* = 0.01 (*E*). Shaded ribbons represent ±1 standard deviation from the average.

Once a non-native tetraploid enters the new mixed-ploidy population, its genetic information can spread by three different non-exclusive pathways; a 4*x* × 4*x* cross, in which case the new genetic information remains at the tetraploid level, and 4*x* × 3*x* or 4*x* × 2*x* cross, which may produce either a tetraploid or a triploid individual. Here, we are primarily concerned with the latter two pathways, as a triploid organism can then produce haploid gametes and backcross with the diploid state inside the population producing a new diploid cytotype. The speed and frequency with which alleles flowing through these pathways enter the diploid state of the new population will then depend on triploid viability and random effects produced by drift (see Methods). We found that the frequency of introgressed alleles per generation as a function of *φ* is qualitatively similar to the effect of *κ*. Even for *φ* = 0.01, the frequency of non-native alleles grows to an average of 20.7% after 20,000 generations (Fig. 3*C*). It should be noted, however, that this effect is in part a consequence of the lower frequency of tetraploid cytotypes in both populations, as described by Eqs. **1** and **2**. By decreasing *φ* and keeping *ν* constant, the frequency of tetraploid cytotypes will decay. Therefore, in *SI Appendix Text S2* we tested the effect of triploid viability by normalizing *ν* as a function of φ (Eq. 2.6), which allowed us to show the same effect decoupled from tetraploid frequencies. Also, in *SI Appendix Text S3*, we further show the evolution of introgression using a deterministic model of a single biallelic locus in the limit *N* ⟶ ∞, where we verify that in the absence of selective pressures, fixation of a non-native neutral allele happens in the span of a few generations.

As each population is fixed for a different allele symmetrical gene flow at the tetraploid level will eventually lead to an equilibrium frequency of 50% of allele types in each population given enough time. Therefore, we were interested in the frequency with which a single mutant allele - subject to drift (see *Methods*) - arising in say, population A, gets established in population B. We initialized our system with both populations fixed for a wild-type variant and after cytotype equilibrium frequencies have been reached, as described by Equation 2.3, we released a mutant neutral allele in population A at the diploid level and measured the number of times the mutant appears in the diploid gene pool of population B or gets lost. We found the probability that the mutant survives and introgresses into the diploid state of population B to be ∼2.3%, with the time it takes to cross the bridge well described by an exponential distribution with an average of ∼43 generations (see *SI Appendix* Fig. S2).

To further understand gene flow through polyploid bridges, we built a fitness map that modulates the probability of successful mating with respect to the genetic distance between individuals and the population’s reference genome, taken to be the optimal (see Methods, equation 1.3). Essentially, the probability that an individual successfully reproduces depends on how well its genetic background approximates an optimal sequence (reference) that represents the environment of its population. We first tested the effect that selection strength, represented by *s*, has on the average frequency of introgressed alleles in each population (Fig. 3D). As expected, the larger the selection strength is, the lower the frequency of non-native alleles that can enter a new gene pool at the diploid level, impeding the system from achieving equilibrium. When genetic divergence is too high, hybrids are subject to strong selective pressures, and introgression of non-native alleles is blocked (Fig. 3*E*). In our simulations, introgression at the diploid level under high genetic divergence between populations is primarily hampered by high selection at the tetraploid state (see *SI Appendix* Fig. S3). Higher recombination rates can, however, increase optimal allele sorting between lineages for introgression down to the diploid states (see *SI Appendix* Fig. S4).

### WGD-mediated adaptive introgression compared to adaptive introgression through diploids

We now turn our attention to the transfer of genes with adaptive value into a new diploid gene pool from a different mixed-ploidy population. WGD-mediated adaptive introgression has been argued to play a substantial role in the adaptation of polyploids for both intracellular and environmental demands (33), but its relevance may extend to diploids by means of the mechanisms explored previously. Since reproductive isolation between diploid species is seldom complete (65), especially in plants (32), the question remains how much gene flow is necessary at the tetraploid level (*κ*) to supersede diploid-mediated adaptive introgression. If diploid states exchange genetic information interspecifically, even if at low rates, we ask how much gene flow at the tetraploid level is required to override diploid-mediated introgression of an adaptive allele into a new diploid gene pool. This question arises naturally when one asks about the relative importance of gene flow between polyploid taxa given that their diploid counterparts are not completely isolated. More specifically, we are interested in the time it takes for an allele with adaptive value to introgress into a new diploid gene pool through polyploid bridges and compare it with interspecific gene flow at the diploid level (Fig. 4).

**Fig. 4.**
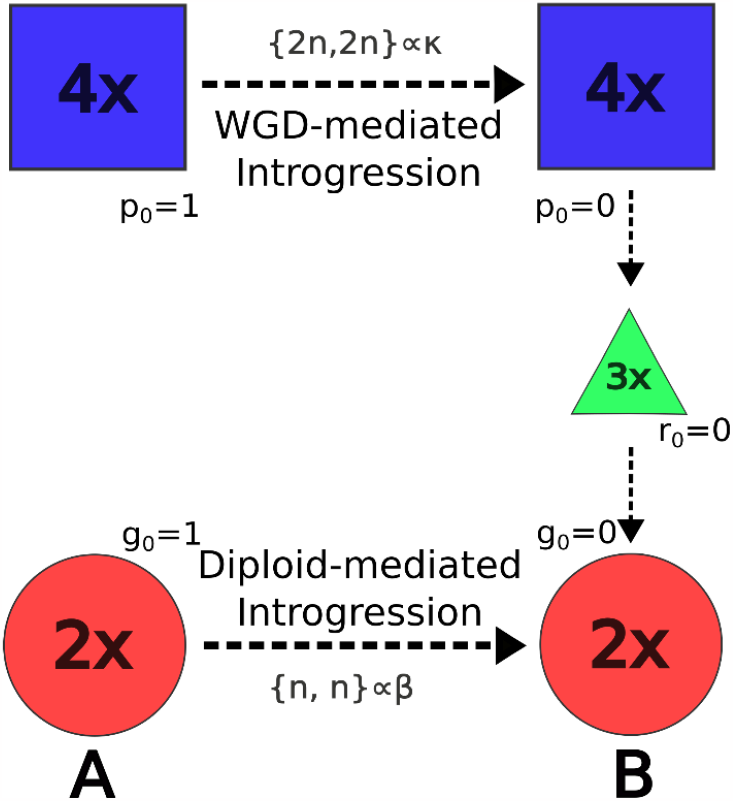
Routes for adaptive introgression in a new diploid gene pool. Two routes are presented for unidirectional gene flows from species A to species B. Species A is initially (*t* = 0) fixed for a beneficial allele whose frequency in the tetraploid (4*x*) and diploid (2*x*) states is given by *p*_*t*_ and *g*_*t*_, respectively. Gene flow from species A to species B is represented by the transfer of gametes (in curly brackets) between gene pools as before. Gene flow at the diploid level occurs with rate *β* and at the tetraploid level with rate *κ*. The speed with which a beneficial allele will introgress in the diploid gene pool of species B (lower left corner) was compared when genetic information flows interspecifically through the route 4*x* ⟶ 4*x* ⟶ 3*x* ⟶ 2*x*, with the case where gene flow occurs through the route 2*x* ⟶ 2*x*, mediated by *β*.

To measure how introgression of a beneficial allele between tetraploids differs from gene flow between diploid states, we built a mainland-island model with unidirectional gene flow from the mainland (species A) into the island (species B) (66) for each ploidy-mediated introgression case. We assumed a single biallelic locus, where the mainland species is fixed for a beneficial allele *A* (both diploids and tetraploids) and the island species is fixed for a deleterious variant *a* (across all cytotypes). The deleterious allele has a negative additive fitness effect *s* on genotypes, whose fitness of the homozygous states is decreased by a factor (1 − *s*). Here, we shall derive the equations that describe adaptive introgression through the route 4*x* (species A) ⟶ 4*x* (species B) ⟶ 3*x* (species B) ⟶ 2*x* (species B), *i*.*e*., polyploidy-mediated introgression. For the tetraploid gene pool in the island there are three possible gametes, which we define as *D*_1_ ≜ *AA, D*_2_ ≜ *Aa* and *D*_3_ ≜ *aa*. Gametes of type *D*_1_ migrate from the tetraploid gene pool at the mainland at a rate κ, which will mix with other diploid gametes produced within the island by tetraploid and triploid genotypes (as described in previous sections). Let *p*_*t*_ and *r*_*t*_ represent the frequency of *A* in the island tetraploid and triploid gene pools, respectively, at time *t*, and *q*_*t*_ = 1 − *p*_*t*_ and *w*_*t*_ = 1 − *r*_*t*_ the frequency of their corresponding deleterious variants. Under random mating and assuming no double reduction during meiosis, with average triploid fertility *φ* = 0.30 (39), we found that the proportions of each diploid gamete type in the island gene pool, following unidirectional gamete migration from the mainland, can be calculated as follows:

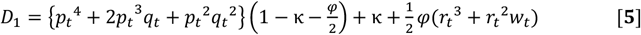

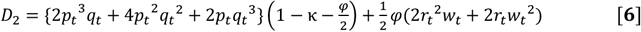

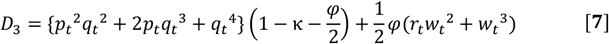

Likewise, we defined haploid gametes in the island as *H*_1_ ≜ *A* and *H*_2_ ≜ *a*, which can be produced by triploid and diploid cytotypes, as in the previous models. Then, if *g*_*t*_ and *z*_*t*_ = 1 − *g*_*t*_ are the frequencies of *A* and *a*, respectively, among the diploid cytotypes, the proportion of haploid gamete types, following unidirectional migration from the mainland diploid species into the island (at rate *β*) and random mating, can be calculated as follows:

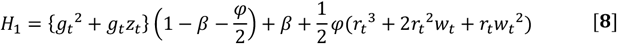

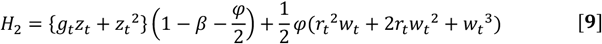

Eqs. **5** to **9** thus define the proportions of all gamete types produced by all cytotypes within the island at each time *t* with ongoing gene flow from the mainland. By computing the gamete types, we can retrieve the genotypes that make up the island population simply by the expansion of the polynomial (*D*_1_ + *D*_2_ + *D*_3_ + *H*_1_ + *H*_2_)^2^, whose frequencies will decay according to additive selection on the deleterious variant. For example, the resulting tetraploid genotype *D*_2_^2^ will have its frequency decreased by a factor (1 − 0.5*s*) as half of its loci are *a*. By collecting all terms of the expansion and computing the frequency of *A* within the diploid gene pool, we find that the recursive equation that describes the evolution of the beneficial mutation among the diploid cytotypes is written as follows:

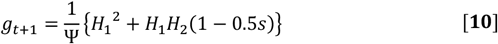

where,

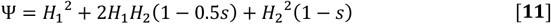

In *SI Appendix Text S4*, we provide a detailed derivation of Eqs. **5** to **9**, prove the result in Equation 3.6, and provide the explicit formulas for *p*_*t*+1_ and *r*_*t*+1_. Notice that when *φ* = 0, the only source of the beneficial mutation in the diploid gene pool comes from diploid-mediated introgression at rate *β* (Eqs. **8** and **9**). When *β* = 0, beneficial mutations only arrive through polyploidy-mediated gene flow (if *φ* > 0), and thus *g*_*t*+1_ is only a function of κ.

Setting *φ* = 0 and *β* = κ, we first measured how the frequency of allele *A* unfolds in time for introgression at the tetraploid level (4*x* ⟶ 4*x, p*_*t*_) and compared it with introgression at the diploid level (2*x* ⟶ 2*x, g*_*t*_) for different values of *s*. That is, for the same introgression rate between tetraploids and diploids, we ask how introgression unfolds between tetraploid and between diploid populations. As expected, the speed with which allele *A* reaches fixation in the receiver populations depends on how strong selection is on the deleterious variant *a*, that is, the higher the selection strength on *a*, the faster introgression occurs (Fig. 5*A*). Additionally, for any selection coefficient *s* > 0, diploid-mediated introgression is always more efficient (*SI Appendix* Fig. S6). Indeed, population genetics theory predicts that deleterious mutations may accumulate faster in polyploids relative to diploids due to a masking effect (67), which has been more recently experimentally confirmed by a number of studies (6, 68). Essentially, due to the higher number of genome copies in the polyploid state, deleterious mutations are masked from selection, and therefore are expected to accumulate in polyploid lineages with much less friction. Alternatively, our results show that this phenomenon decreases the efficiency with which the introgression of a beneficial allele occurs in a tetraploid as compared to a diploid gene pool.

**Fig. 5.**
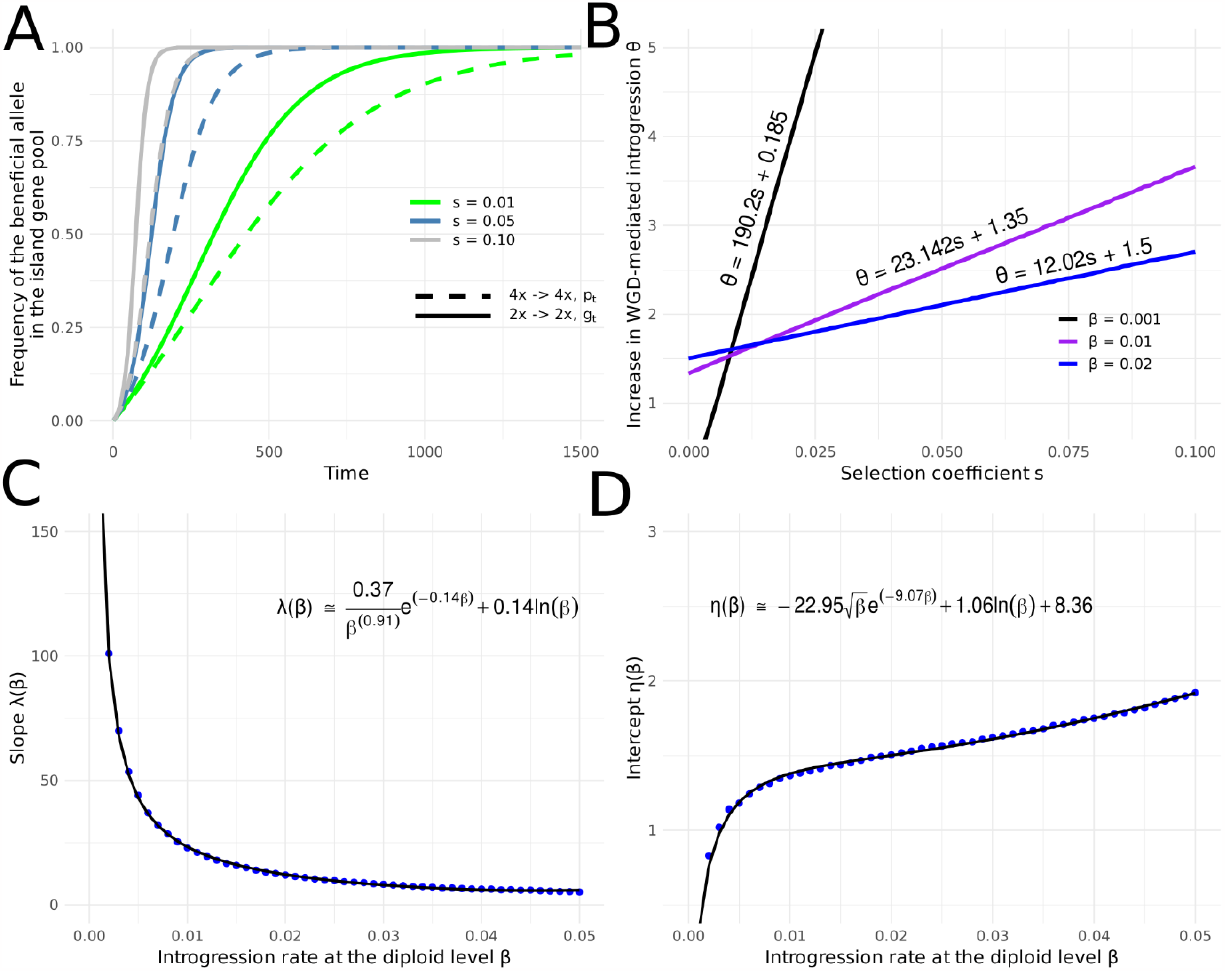
Adaptive introgression efficiency comparison between gene flow mediated by diploids and tetraploids in the mainland-island model. (A) Frequency of the beneficial allele *A* as a function of time (iterations) and selection strength *s* in the island at the diploid level through unidirectional gene flow from the mainland’s diploid population (*g*_*t*_, solid lines), and at the tetraploid level through unidirectional gene flow from the mainland’s tetraploid population (*p*_*t*_, dashed lines). Gene flow rates are the same in both cases: *β* = *α* = 10^−3^. Additionally, *φ* = 0. (B) Minimum increase in WGD-mediated introgression rate *θ* to achieve fixation of the beneficial allele within the same time interval as achieved by diploid-mediated introgression (*φ* = 0.30). (C and D) Relationship between the slopes and intercepts of the linear equations (B) as a function of introgression at the diploid level *β*. Equation coefficients were retrieved by the Levenberg-Marquadt algorithm.

To understand how both two routes (4*x* ⟶ 4*x* ⟶ 3*x* ⟶ 2*x* and 2*x* ⟶ 2*x*) differ for an introgressing beneficial allele, we measured the time with which the beneficial mutation gets fixed within the diploid gene pool through diploid-mediated gene flow for low introgression rates (*β* ≪ 1), and computed the WGD-mediated introgression rate (κ_*min*_) for which the allele gets fixed in the same diploid pool within the same time interval (see *SI Appendix Algorithm S2* for a description of the algorithm). We then computed the factor *θ* = κ_*min*_/*β* as it more clearly informs us on the minimum increase in WGD-mediated introgression rate necessary for introgression through polyploidy to be at least as efficient as diploid-mediated adaptive introgression. We found that *θ* is a linear function of the selection coefficient *s* (Fig. 5*B*), that is, for each unit increase in *s* the amount of WGD-mediated gene flow to override diploid-mediated introgression increases linearly. However, the degree of the increase (slope) as well as the minimum increase for a neutral allele (intercept when *s* = 0) has a complex relationship with the introgression rate at the diploid level (*β*). Thus, in general, we can write *θ* as a function of *β* as follows:

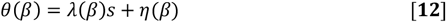

That is, *θ* is a linear function of the selection coefficient *s*, where the slope *λ*(*β*) and intercept *η*(*β*) are unkown functions of *β*. To find these unkown functions we modelled separately the slope and intercept with respect to the introgression rate at the diploid level. Both the slope *λ*(*β*) (Fig. 5*C*) and intercept *η*(*β*) (Fig. 5D) were found to be non-trivial functions of *β*, whose forms we could only approximate locally (0.001 ≤ *β* ≤ 0.05). These approximations allowed us to retrieve a general equation to estimate the minimum amount of gene flow at the tetraploid level necessary to override adaptive introgression mediated by diploid states. Realizing κ_*min*_ = *θβ*, we write:

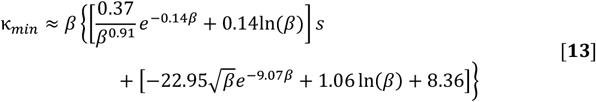

For any *s*, Eq. **13** tells us that any κ > 0 is sufficient to allow WGD-mediated introgression when diploids are isolated (*β* ⟶ 0) consistent with our arguments in the previous sections. The use of Eq. **13** is subject to two experimental challenges in natural or artificial systems. First, the introgression rate at the diploid level *β* must be recovered, which can be determined by analyses of gene flow using molecular markers (69), for example. A second challenge concerns the determination of the selection coefficient *s* that a given allele is subject to relative to an adaptive variant present in the population from which gene flow is being measured. However, a first lower-bound to gene flow at the tetraploid level necessary to overrride gene flow at the diploid level can be achieved by setting *s* = 0, which informs the experimentalist on the amount of gene flow between tetraploids necessary for polyploid bridges to display a relevant impact on the distribution of neutral genetic material across species boundaries.

## DISCUSSION

Understanding how genetic material flows within ecosystems is crucial for our understanding of biological evolution. By generalizing the well-known model of Felber (48) for cytotype dynamics, we show that the transfer of genetic information between different diploid species is possible under the current understanding of interploidy mating dynamics and WGD-potentiated hybridization. A notable example of introgression, potentially explained by gene flow through polyploid bridges, pertains to genes encoding the *C*_4_ phosynthetic pathway in *Alloteropsis semialata*, which have been laterally acquired from the polyploid *Setaria palmifolia* complex, most likely in tropical Africa, where both species co-occur (42). Most of the non-*C*_4_ individuals in *A. semialata* are diploids, with *C*_4_ individuals ranging from diploid to dodecaploid levels (41, 70). Although triploid hybrids have not been reported so far for *A. semialata*, the most plausible path through which genetic information coding for *C*_4_ biochemistry could have introgressed into the diploid level of the species would be through compatible crosses that required the presence of a triploid intermediate, from which a cross of the type 3*x* × 2*x* would allow the information to spread down to the diploid level. An alternative hypothesis may relate the presence of *C*_4_ diploid individuals to the process of diploidization. However, despite the fact that diploidization is a slow process measured at macroevolutionary scales (11), there is direct experimental evidence that crosses between photosynthetic types in *A. semialata* are viable (70). That is, following introgression from polyploid *S. palmifolia*, information coding for *C*_4_ biochemistry had to necessarily travel through the different ploidy levels in *A. semialata* to reach the diploid level, where *C*_4_ and non-*C*_4_ individuals are then capable of exchanging genetic material.

It should be noted that the extent to which WGD facilitates interspecific introgression is not entirely clear (33), which constrains our ability to unequivocally ascertain the impact of polyploid bridges on community-level evolutionary dynamics. For instance, in a recent study, Marburger et al. (27) were able to identify bidirectional introgression of genes controlling meiotic processes between tetraploid cytotypes of *A. arenosa* and *A. lyrata*. These shared introgressed genes, implicated in mechanisms like meiotic double strand break formation, were deemed instrumental for the emergence of stable meiotic processes in both lineages. Nevertheless, the variability in meiotic stability found in the studied populations suggests that interspecific gene flow at the polyploid level of both species depends on optimal sorting of segregating alleles for stable lineages to emerge, which is supported by signatures of selective sweep across their genomes (27). Here, we demonstrated that although WGD-mediated interspecific introgression down to the diploid level is constrained with surging genetic divergence between mixed-ploidy populations, increased recombination rates can relieve selection pressures, thereby facilitating the influx of new allele variants into a new gene pool. Interestingly, polyploidization has also been shown to increase recombination rates in *Arabidopsis* (71), yet a clear understanding of the relationship between genetic divergence and WGD-mediated gene flow remains elusive.

While genetic incompatibility is one of the mechanisms of reproductive isolation that is potentially circumvented by polyploidy, WGD-potentiated hybridization may also favor the transfer of genetic material across diverging populations by evading pre-zygotic barriers. Indeed, since Levin (56) postulated the principle of minority cytotype exclusion, the hypothesis that ecological disparities must exist between ploidy levels for successful establishment of polyploid cytotypes, has led researchers to identify a number of cases where niche differentiation or expansion following WGD occurs (15, 16, 28, 72-74). In particular, niche expansion of polyploid organisms relative to their diploid ancestors might promote secondary contact among allopatric populations of a specific taxon, restoring therefore gene flow and counteracting reproductive isolation (33). Within this paradigm, the interpretation of the migration coefficient used in our model (*κ*) could span both evolutionary and ecological dimensions. On the one hand, WGD may directly breach species barriers (28, 75) by overcoming putative post-zygotic limitations, and on the other hand promote intraspecific gene flow among diverging populations via secondary contact or niche overlaps, as possibly observed in *Senecius carniolicus* (76).

For the purposes of our theoretical investigation, we neglected the possibility of triploid gamete formation by triploid cytotypes. It has been shown that the frequency of triploid gametes is on average higher than the sum of diploid and haploid gametes produced by triploids (39). This will lead to an increased probability of tetraploid formation relative to the present study and would hence increase gene flow from diploid to tetraploids relative to gene flow from tetraploids to diploids (see *SI Appendix Text S5*). Despite the bias in gene flow due to differential production of gametes in triploids, the path for interploidy bidirectional flux of genetic material should remain open. This contrasts with Cao et al.’s recent assertion (77) that hybrid triploid self-fertilization invariably yields tetraploids, correlating with a reproductive barrier thwarting gene flow from tetraploids back to diploid levels, termed a “triploid block”. Therefore, in *SI Appendix Text S5*, we further show that this effect is also a trivial consequence of the probabilities for merging different gamete ploidy levels during self-fertilization in triploid cytotypes. The model presented here is anchored in our current understanding of *interploidy* mating dynamics, and self-fertilization - though potentially important in the establishment of new cytotypes - is tangential to this study’s analytical purview.

Throughout this paper, we assumed additive fitness effects that are proportional to the ploidy level. However, the hybridization of two polyploid species, or allopolyploidization, may result in nonadditive gene expression patterns on hybrids (78), which may influence fitness at the polyploid level and therefore impact the efficiency with which genetic material diffuses through ploidy levels in the recipient gene pool. The extent to which the emergence of polyploid bridges influences adaptation and disturb allele frequencies in real biological systems is yet to be determined. The flux of genes among diploid states from different species may depend on the rate at which genes flow among polyploids, which in turn may be conditioned on the spatial and temporal sequences with which higher-ploidy species come into contact. Furthermore, the fitness map used in this study imposes selection in an exogeneous manner, that is, the fitness of an organism depends on the distance of its genome to an external sequence that represents the optimum of the environment in which the individual inhabits. Selection on hybrids may as well be based on disruption of parental gene combinations, emerging therefore as the product of negative epistatic interactions. For instance, non-Mendelian segregation patterns on synthetic hybrids between polyploids from cotton have been attributed to epistatic interactions following the merging of parental genomes, whose coadapted genes constrain the genomic composition of hybrid offspring (79). The findings portrayed in the present study can serve as a guide for further theoretical investigation on the phenomenon, as well as spur experimental research to potentially uncover new mechanisms of gene flow within ecosystems.

While polyploidy may be a transient phenomenon, i.e., an evolutionary dead-end (3), its periodic emergence in the evolutionary history of life can serve as a bridge to connect diverged taxa and redistribute genetic information within ecosystems. Here, we have provided a systematic study on the cytotype dynamics of mixed-ploidy populations and showed that, when genetic material is explicitly accounted for, WGD may promote lateral gene transfer. Further investigations on the significance of polyploid bridges might provide insightful results into the delicate interactions that weave biological networks through evolutionary history.

## METHODS

Our stochastic individual-based simulation contains a population of size *N*, where each discrete diploid individual is represented by a set of 2 vectors, called chromosomes, with *L* elements. Chromosome elements are taken from the set {0, 1}, representing biallelic loci. In the standard model, all individuals are identical, i.e., *S*_*i,k*_ = *S*_*j,k*_, ∀ *i, j* ∈ {*N*} and *k* ∈ {*L*}. Mating is synchronous and is performed by randomly choosing two individuals, with replacement, from their corresponding mixed-ploidy population, whose gametes formed (see below) are then joined to make up the genome of the offspring, until the carrying capacity *N* is restored. Notice that the probability that an individual is not chosen to reproduce in each generation is (1 − 2/*N*)^*N*^, which approaches *e*^−2^ in the limit *N* ⟶ ∞, corresponding to drift. When mating is completed then bidirectional migration occurs at the tetraploid level where each individual has a probability *κ* of entering the next population. Each iteration, or generation, corresponds to a repetition of this process for a total of 20,000 generations.

A mating event consists of randomly selecting two individuals from the same population, with each one undergoing meiosis, where recombination among *rL* randomly selected loci (*r* ≪ 1) takes place. In a diploid individual, meiosis pairs the two chromosomes and recombination occurs. Then, a gamete is one of the resulting chromosomes chosen with equal probability, or both chromosomes, *i*.*e*., unreduced diploid gametes, with probability *ν*. In triploids, recombination occurs randomly between two of the three available chromosomes mimicking chromosome pairing during triploid meiosis as shown in previous experimental studies (80). In triploid meiosis three outcomes are possible: i) the individual produces no gametes with probability (1 − φ) owing to differential contribution to the gene pool due to viability, in which case the mating event fails, ii) a haploid gamete, or iii) a diploid gamete is randomly chosen out of the three available chromosomes. We set outcomes ii) and iii) to happen with equal probability (see Eqs. **1** and **2** in Results for a mathematical description). Finally, meiosis in a tetraploid individual results in the random formation of two bivalents out of the four available chromosomes, with no preferential chromosome pairing, and a gamete is one of the bivalent pairs chosen with equal probability after recombination has occurred, following previous work (20). Although chromosome pairing in polyploid meiosis can take up different forms depending on polyploid origin (54), our goal is to isolate the effect of recombination, and not to provide a comprehensive mechanistic understanding of the possible results following differential recombination patterns in polyploid meiosis. In *SI Appendix* Fig. S7, the reader will find an illustration detailing the meiosis procedure used in this work.

To understand the dynamics of the model under a selection regime, we developed a fitness map, assuming allele codominance, that relates the genomes of individuals to a sequence that represents the optimum phenotype (a reference genome randomly defined) in their respective mixed-ploidy populations. Let *d*(*G*_*i*_; *G*_*D*_) be the total distance between the genome of individual *i* to its respective population’s genome *D* (a single vector), relative to chromosome length *L*. Then, for each loci *k* in each chromosome *p* of individual *i* we define the distance to be:

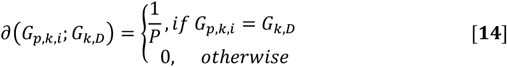

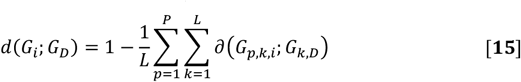

where *P* is the ploidy level of individual *i*.

For a mating trial to be successful we require that both individuals have a fitness value larger than a random number generated with uniform distribution in the interval (0,1), otherwise the system will choose other pair of individuals from the same deme until a successful mating event occurs. The fitness *f*of an individual is given by an exponentially decaying function of the form:

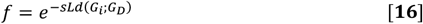

where *s* represents the selection strength that the populations shall be subject to. The genetic distance between demes is assigned upon initialization of the simulation, where the number of alleles that can differ between demes’ sequences are randomly selected to match the desired genetic divergence (see *SI Appendix Algorithm S1* for implementation strategies).

### Data, Materials, and Software Availability

Documentation and software to generate the results presented in this work will be made available upon publication of the manuscript. All additional data and pseudo-codes relevant for this study are available in *SI Appendix*.

## ACKNOWLEDGMENTS

We thank Arthur Zwaenepoel (Unité Évolution, Écologie, Paléontologie, Université de Lille 1, Villeneuve d’Ascq, France) for helpful discussions. This work was supported by the European Research Council under the European Union’s Horizon 2020 Research and Innovation program (No. 833522) and from Ghent University (Methusalem funding, BOF.MET.2021.0005.01) (to Y.V.d.P.).

## Author contributions

F.K., Q.B., F.M., M.V.M., D.B., and Y.V.d.P. designed research; F.K., Q.B., and F.M. performed research; F.K., Q.B., F.M., D.B., Y.V.d.P analyzed data; and F.K. and Y.V.d.P. wrote the paper.

## References

1. S. De Bodt, S. Maere, Y. Van de Peer, Genome duplication and the origin of angiosperms. Trends Ecol. Evol. 20, 591–597 (2005).

2. D. E. Soltis, J. G. Burleigh, Surviving the K-T mass extinction: new perspectives of polyploidization in angiosperms. Proc. Natl. Acad. Sci. U.S.A. 106, 5455–5456 (2009).

3. Y. Van de Peer, E. Mizrachi, K. Marchal, The evolutionary significance of polyploidy. Nat. Rev. Genet. 18, 411–424 (2017).

4. Y. Van de Peer, T. L. Ashman, P. S. Soltis, D. E. Soltis, Polyploidy: an evolutionary and ecological force in stressful times. Plant Cell 33, 11–26 (2021).

5. S. P. Otto, The evolutionary consequences of polyploidy. Cell 131, 452–462 (2007).

6. P. Monnahan et al., Pervasive population genomic consequences of genome duplication in Arabidopsis arenosa. Nat. Ecol. Evol. 3, 457–468 (2019).

7. G. Blanc, K. H. Wolfe, Widespread paleopolyploidy in model plant species inferred from age distributions of duplicate genes. Plant Cell 16, 1667–1678 (2004).

8. A. Zwaenepoel, Y. Van de Peer, inference of ancient whole-genome duplications and the evolution of gene duplication and loss rates. Mol. Biol. Evol. 36, 1384–1404 (2019).

9. L. Cai et al., Widespread ancient whole-genome duplications in Malpighiales coincide with Eocene global climatic upheaval. New Phytol. 221, 565–576 (2019).

10. P. Y. Novikova, N. Hohmann, Y. Van de Peer, Polyploid Arabidopsis species originated around recent glaciation maxima. Curr. Opin. Plant Biol. 42, 8–15 (2018).

11. A. K. Redmond, D. Casey, M. K. Gundappa, D. J. Macqueen, A. McLysaght, Independent rediploidization masks shared whole genome duplication in the sturgeon-paddlefish ancestor. Nat. Commun. 14, 2879 (2023).

12. K. Vanneste, G. Baele, S. Maere, Y. Van de Peer, Analysis of 41 plant genomes supports a wave of successful genome duplications in association with the Cretaceous-Paleogene boundary. Genome Res. 24, 1334–1347 (2014).

13. D. A. Levin, D. E. Soltis, Factors promoting polyploid persistence and diversification and limiting diploid speciation during the K-Pg interlude. Curr Opin Plant Biol 42, 1–7 (2018).

14. Q. Bafort, T. Wu, A. Natran, O. De Clerck, Y. Van de Peer, The immediate effects of polyploidization of Spirodela polyrhizachange in a strain-specific way along environmental gradients. Evol. Lett. 7, 37–47 (2023).

15. A. Rice et al., The global biogeography of polyploid plants. Nat. Ecol. Evol. 3, 265–273 (2019).

16. K. T. David, Global gradients in the distribution of animal polyploids. Proc. Natl. Acad. Sci. U.S.A. 119, e2214070119 (2022).

17. M. Ebadi et al., The duplication of genomes and genetic networks and its potential for evolutionary adaptation and survival during environmental turmoil. Proc. Natl. Acad. Sci. U.S.A. 120, e2307289120 (2023).

18. I. Mayrose et al., Recently formed polyploid plants diversify at lower rates. Science 333, 1257 (2011).

19. D. A. Levin, Why polyploid exceptionalism is not accompanied by reduced extinction rates. Plant Syst. Evol. 305, 1–11 (2019).

20. F. Kauai, F. Mortier, S. Milosavljevic, Y. Van de Peer, D. Bonte, Neutral processes underlying the macro eco-evolutionary dynamics of mixed-ploidy systems. Proc. Biol. Sci. 290, 20222456 (2023).

21. Y. Wang et al., An ancient whole-genome duplication event and its contribution to flavor compounds in the tea plant (Camellia sinensis). Hort. Res. 8, 176 (2021).

22. J. W. Clark, Genome evolution in plants and the origins of innovation. New Phytol. n/a (2023). doi: 10.1111/nph.19242

23. M. Xiao et al., Seagrass genomes reveal a hexaploid ancestry facilitating adaptation to the marine environment. Nat. Plants (2023). In press.

24. W. Bleeker, Hybridization and Rorippa austriaca (Brassicaceae) invasion in Germany. Mol. Ecol. 12, 1831–1841 (2003).

25. M. H. Jorgensen, D. Ehrich, R. Schmickl, M. A. Koch, A. K. Brysting, Interspecific and interploidal gene flow in Central European Arabidopsis (Brassicaceae). BMC Evol. Biol. 11, 346 (2011).

26. B. J. Arnold et al., Borrowed alleles and convergence in serpentine adaptation. Proc. Natl. Acad. Sci. U.S.A. 113, 8320–8325 (2016).

27. S. Marburger et al., Interspecific introgression mediates adaptation to whole genome duplication. Nat. Commun. 10, 5218 (2019).

28. P. Y. Novikova et al., Polyploidy breaks speciation barriers in Australian burrowing frogs Neobatrachus. PLoS Genet. 16, e1008769 (2020).

29. D. Ståhlberg, M. Hedrén, Evolutionary history of the Dactylorhiza maculata polyploid complex (Orchidaceae). Biol. J. Linn. Soc. Lond. 101, 503–525 (2010).

30. P. A. Christin et al., Multiple photosynthetic transitions, polyploidy, and lateral gene transfer in the grass subtribe Neurachninae. J. Exp. Bot. 63, 6297–6308 (2012).

31. C. Lafon-Placette et al., Endosperm-based hybridization barriers explain the pattern of gene flow between Arabidopsis lyrata and Arabidopsis arenosa in Central Europe. Proc. Natl. Acad. Sci. U.S.A. 114, E1027–E1035 (2017).

32. J. Mallet, Hybrid speciation. Nature 446, 279–283 (2007).

33. R. Schmickl, L. Yant, Adaptive introgression: how polyploidy reshapes gene flow landscapes. New Phytol. 230, 457–461 (2021).

34. W. Albertin, P. Marullo, Polyploidy in fungi: evolution after whole-genome duplication. Proc. Biol. Sci. 279, 2497–2509 (2012).

35. J. Steensels, B. Gallone, K. J. Verstrepen, Interspecific hybridization as a driver of fungal evolution and adaptation. Nat. Rev. Microbiol. 19, 485–500 (2021).

36. E. L. Breese, E. J. Lewis, G. M. Evans, Interspecies hybrids and polyploidy. Philos. Trans. R. Soc. Lond. B Biol. Sci. 292, 487–497 (1981).

37. J. Majka et al., Cytogenetic and molecular genotyping in the allotetraploid Festuca pratensis × Lolium perenne hybrids. BMC Genomics 20, 367 (2019).

38. B. Arnold, S. T. Kim, K. Bomblies, single geographic origin of a widespread autotetraploid Arabidopsis arenosa lineage followed by interploidy admixture. Mol. Biol. Evol. 32, 1382–1395 (2015).

39. J. Ramsey, D. W. Schemske Pathways, mechanisms, and rates of polyploid formation in flowering plants. Annu. Rev. Ecol. Syst. 29, 467–501 (1998).

40. T. L. Burton, B. C. Husband, Fecundity and offspring ploidy in matings among diploid, triploid and tetraploid Chamerion angustifolium (Onagraceae): consequences for tetraploid establishment. Heredity 87, 573–582 (2001).

41. J. K. Olofsson et al., Low dispersal and ploidy differences in a grass maintain photosynthetic diversity despite gene flow and habitat overlap. Mol. Ecol. 30, 2116–2130 (2021).

42. P.-A. Christin et al., Adaptive evolution of C4 photosynthesis through recurrent lateral gene transfer. Curr. Biol. 22, 445–449 (2012).

43. C. Phansopa, L. T. Dunning, J. D. Reid, P. A. Christin, lateral gene transfer acts as an evolutionary shortcut to efficient C4 biochemistry. Mol. Biol. Evol. 37, 3094–3104 (2020).

44. G. P. Tiley et al., Estimation of species divergence times in presence of cross-species gene flow. Syst. Biol. 72, 820–836 (2023).

45. M. Slatkin, Gene flow in natural populations. Ann. Rev. Ecol. Syst. 16, 393–430 (1985).

46. J. P. Spoelhof, R. Keeffe, S. F. McDaniel, Does reproductive assurance explain the incidence of polyploidy in plants and animals? New Phytol.t 227, 14–21 (2020).

47. S. P. Otto, J. Whitton, Polyploid incidence and evolution. Annu. Rev. Genet. 34, 401–437 (2000).

48. F. Felber, Establishment of a tetraploid cytotype in a diploid population: Effect of relative fitness of the cytotypes. J. Evol. Biol. 4, 195–207 (1991).

49. F. Felber, J. D. Bever, Effect of triploid fitness on the coexistence of diploids and tetraploids. Biol. J. Linn. Soc. Lond. 60, 95–106 (1997).

50. B. C. Husband, The role of triploid hybrids in the evolutionary dynamics of mixed-ploidy populations. Biol. J. Linn. Soc. Lond. 82, 537–546 (2004).

51. W. E. Van Drunen, J. Friedman, Autopolyploid establishment depends on life-history strategy and the mating outcomes of clonal architecture. Evolution 76, 1953–1970 (2022).

52. E. J. Baack, Ecological factors influencing tetraploid establishment in snow buttercups (Ranunculus adoneus, Ranunculaceae): minority cytotype exclusion and barriers to triploid formation. Am. J. Bot. 92, 1827–1835 (2005).

53. T. Peckert, J. Chrtek, Mating Interactions between coexisting diploid, triploid and tetraploid cytotypes of Hieracium echioides (Asteraceae). Folia Geobot. 41, 323–334 (2006).

54. M. Cifuentes, L. Grandont, G. Moore, A. M. Chevre, E. Jenczewski, Genetic regulation of meiosis in polyploid species: new insights into an old question. New Phytol. 186, 29–36 (2010).

55. J. M. Kreiner, P. Kron, B. C. Husband, Evolutionary dynamics of unreduced gametes. Trends Genet. 33, 583–593 (2017).

56. D. A. Levin, Minority cytotype exclusion in local plant populations. Taxon 24, 35–43 (1975).

57. C. C. F. Schinkel et al., Pathways to polyploidy: indications of a female triploid bridge in the alpine species Ranunculus kuepferi (Ranunculaceae). Plant Syst. Evol. 303, 1093–1108 (2017).

58. M. M. Li et al., Triploid cultivars of Cymbidium act as a bridge in the formation of polyploid plants. Front. Plant Sci. 13, 1029915 (2022).

59. Æ. T. Thórsson, S. Pálsson, M. Lascoux, K. Anamthawat-Jónsson, Introgression and phylogeography of Betula nana (diploid), B. pubescens (tetraploid) and their triploid hybrids in Iceland inferred from cpDNA haplotype variation. J. Biogeogr. 37, 2098–2110 (2010).

60. Æ. T. Thórsson, E. Salmela, K. Anamthawat-Jonsson, Morphological, cytogenetic, and molecular evidence for introgressive hybridization in birch. J. Hered. 92, 404–408 (2001).

61. I. M. Henry et al., Aneuploidy and genetic variation in the Arabidopsis thaliana triploid response. Genetics 170, 1979–1988 (2005).

62. P. Farhat et al., Gene flow between diploid and tetraploid junipers - two contrasting evolutionary pathways in two Juniperus populations. BMC Evol. Biol. 20, 148 (2020).

63. D. Ståhlberg, Habitat differentiation, hybridization and gene flow patterns in mixed populations of diploid and autotetraploid Dactylorhiza maculata s.l. (Orchidaceae). Evol. Ecol. 23, 295–328 (2007).

64. Q. Qin et al., Abnormal chromosome behavior during meiosis in the allotetraploid of Carassius auratus red var. (♀) × Megalobrama amblycephala (♂). BMC Genet. 15, 95 (2014).

65. A. M. Westram, S. Stankowski, P. Surendranadh, N. Barton, What is reproductive isolation? J. Evol. Biol. 35, 1143–1164 (2022).

66. B. Reinhard, A survey of migration-selection models in population genetics. Discrete Continuous Dyn. Syst. Ser. B 19IS - 4SP - 883, 959 (2014).

67. R. R. Hill, Jr., Selection in autotetraploids. Theor. Appl. Genet. 41, 181–186 (1970).

68. J. L. Conover, J. F. Wendel, Deleterious mutations accumulate faster in allopolyploid than diploid cotton (Gossypium) and unequally between subgenomes. Mol. Biol. Evol. 39, msac024 (2022).

69. D. M. Arias, L. H. Rieseberg, Gene flow between cultivated and wild sunflowers. Theor. Appl. Genet. 89, 655–660 (1994).

70. M. E. Bianconi et al., Contrasted histories of organelle and nuclear genomes underlying physiological diversification in a grass species. Proc. R. Soc. Lond. B Biol. Sci. 287, 20201960 (2020).

71. A. Pecinka, W. Fang, M. Rehmsmeier, A. A. Levy, O. Mittelsten Scheid, Polyploidization increases meiotic recombination frequency in Arabidopsis. BMC Biol. 9, 24 (2011).

72. A. E. Baniaga, H. E. Marx, N. Arrigo, M. S. Barker, Polyploid plants have faster rates of multivariate niche differentiation than their diploid relatives. Ecol. Lett. 23, 68–78 (2020).

73. Y. F. Molina-Henao, R. Hopkins, Autopolyploid lineage shows climatic niche expansion but not divergence in Arabidopsis arenosa. Am. J. Bot. 106, 61–70 (2019).

74. N. Padilla-García et al., The importance of considering the evolutionary history of polyploids when assessing climatic niche evolution. J. Biogeogr. 50, 86–100 (2023).

75. I. Rešetnik, P. Schönswetter, M. Temunović, M. H. J. Barfuss, B. Frajman, Diploid chastity vs. polyploid promiscuity – Extensive gene flow among polyploid cytotypes blurs genetic, morphological and taxonomic boundaries among Dinaric taxa of Knautia (Caprifoliaceae). Perspect. Plant Ecol. Evol. Syst. 59, 125730 (2023).

76. M. Sonnleitner et al., Ecological differentiation of diploid and polyploid cytotypes of Senecio carniolicus sensu lato (Asteraceae) is stronger in areas of sympatry. Ann. Bot. 117, 269–276 (2016).

77. Y. Cao et al., Genome balance and dosage effect drive allopolyploid formation in Brassica. Proc. Natl. Acad. Sci. U.S.A. 120, e2217672120 (2023).

78. S. Bottani, N. R. Zabet, J. F. Wendel, R. A. Veitia, Gene expression dominance in allopolyploids: hypotheses and models. Trends Plant Sci. 23, 393–402 (2018).

79. C. X. Jiang et al., Multilocus interactions restrict gene introgression in interspecific populations of polyploid Gossypium (cotton). Evolution 54, 798–814 (2000).

80. M. L. Becak, W. Becak, Further studies on polyploid amphibians (Ceratophrydidae). 3. Meiotic aspects of the interspecific triploid hybrid: Odontophrynus cultripes (2n=22) x O. americanus (4n=44). Chromosoma 31, 377–385 (1970).

