## Supporting Information for "Interspecific transfer of genetic information through polyploid bridges"

#### Text S1. Derivation of equilibrium frequencies and bifurcation analysis

As discussed in the main manuscript, in a panmictic unit, with unreduced gametes frequency $\upsilon$ and triploid fertility φ, after one generation of random mating, the frequency of new gamete types, i.e., haploids $g_1{}'$ and diploids $g_2{}'$ as a function of the previous generation's gametes $g_1$ and $g_2$ are given by:

$$g_1{}' = \frac{g_1{}^2(1-\upsilon) + \varphi g_1 g_2}{g_1{}^2 + 2\varphi g_1 g_2 + g_2{}^2} \qquad \text{S1.1}$$

$$g_2{}' = \frac{\upsilon g_1{}^2 + \varphi g_1 g_2 + g_2{}^2}{g_1{}^2 + 2\varphi g_1 g_2 + g_2{}^2} \qquad \text{S1.2}$$

For simplicity, since $g_2 = 1 - g_1$, we solve for the equilibrium frequencies by setting $g_1{}' = g_1 = g_1{}^*$ as follows:

$$g_1{}^* = \frac{g_1{}^{*\,2}(1-\upsilon) + \varphi g_1{}^*(1-g_1{}^*)}{g_1{}^{*\,2} + 2\varphi g_1{}^*(1-g_1{}^*) + (1-g_1{}^*)^2} \qquad \text{S1.3}$$

Dividing both sides of Eq. $S1.3$ by $g_1{}^*$ and moving all terms to the RHS we get:

$$0 = g_1{}^*(1-\upsilon) + \varphi(1-g_1{}^*) - g_1{}^{*\,2} - 2\varphi g_1{}^*(1-g_1{}^*) - (1-g_1{}^*)^2 \qquad \text{S1.4}$$

After expanding Eq. $S1.4$ and factoring out all terms with equal power we obtain the parabola:

$$0 = 2g_1{}^{*\,2}(\varphi - 1) + g_1{}^*(3 - \upsilon - 3\varphi) + (\varphi - 1) \qquad \text{S1.5}$$

whose roots are given by the quadratic formula:

$$g_1^* = \frac{3\varphi + \upsilon - 3}{4(\varphi - 1)} \pm \sqrt{\frac{(3 - 3\varphi - \upsilon)^2}{16(\varphi - 1)^2} - \frac{1}{2}} \qquad \text{S1.6}$$

Equation $S1.5$ yields a concave down parabola with two equilibrium points; one unstable and one stable (see *SI Appendix* Fig. S8). Notice that because by construction our model always starts with a population consisting of only diploids, only haploid gametes are allowed at time zero and therefore the system will always converge to the first equilibrium encountered which is the stable equilibrium (right intersection on the x-axis). A saddle-node bifurcation occurs when both equilibrium points coalesce into a half-stable equilibrium point. Following even higher φ and $\upsilon$, the parabola is dragged below the x-axis (*SI Appendix* Fig. S8), and thus the system converges to an equilibrium where the frequency of haploid gametes is zero, i.e., tetraploids take over. To find the bifurcation point, $\bar{\upsilon}$, we simply take the first derivative of Eq. $S1.5$ with respect to ${g_1}^*$ and solve the following system of equations for $\upsilon$:

$$\begin{cases} 2{g_1}^{*\,2}(\varphi - 1) + {g_1}^*(3 - \upsilon - 3\varphi) + (\varphi - 1) = 0 \\ 4{g_1}^*(\varphi - 1) + (3 - \upsilon - 3\varphi) = 0 \end{cases} \qquad \text{S1.7}$$

By solving for $(\varphi - 1)$ in the second equation and plugging the result into the first equation we get:

$$g_1 = \frac{1}{\sqrt{2}} \qquad \text{S1.8}$$

Substituting $S1.8$ back into the first equation of $S1.7$ and solving for $\upsilon$, we conclude that:

$$\bar{\upsilon} = (1 - \varphi)(2\sqrt{2} - 3) \qquad \text{S1.8}$$

which verifies the result in Eq. $4$ in the main manuscript.

#### Text S2. Normalization of unreduced gametes frequency to compare triploid fertility decoupled from cytotype frequencies

Here, we wish to normalize $\upsilon$ as a function of $\varphi$. The normalization is necessary because, as described in the first Section of the main manuscript, as we change $\varphi$ the frequency of cytotypes is disturbed to new equilibrium points. Then, the impact of $\varphi$ on the frequency of introgressed alleles will be masked by changes in the frequency of tetraploid cytotypes in the population, which in turn modulates the rate with which alleles are transported between demes for a given value of $\kappa$. As shown in the main manuscript, equations $2.1 - 2.5$ converge to stable equilibria $x_2^* = 0.665$, $x_3^* = 0.301$ and $x_4^* = 0.034$ for $\upsilon = 0.10$ and $\varphi = 0.30$, given by numerical solutions of the iterated maps. To find $\upsilon$ for any given $\varphi$, such that the same equilibria are obtained, we can use one of the equations in $2.3 - 2.5$ from the main manuscript. For simplicity, let us rewrite equation 2.3 in equilibrium as follows:

$$x_2^* = \frac{[x_2^*(1-\upsilon) + 0.5x_3^*\varphi]^2}{[x_2^* + x_3^*\varphi + x_4^*]^2} \qquad \text{S2.1}$$

where $[x_2^* + x_3^*\varphi + x_4^*]^2 = \Omega$ in the main manuscript. After some algebraic manipulation we arrive at:

$$0 = x_2^*\upsilon^2 - (2x_2^* + x_3^*\varphi)\upsilon + \psi \qquad \text{S2.2}$$

with,

$$\psi = x_2^* + x_3^*\varphi + \frac{{x_3^*}^2\varphi^2}{4x_2^*} - \Omega \qquad \text{S2.3}$$

From S3.3 and S2.3 we find $\upsilon$ for any $\varphi$ by simply computing the roots of S3.2:

$$\upsilon = \frac{(2x_2^* + x_3^*\varphi) \pm \sqrt{[-(2x_2^* + x_3^*\varphi)]^2 - 4x_2^*\psi}}{2x_2^*} \qquad \text{S2.4}$$

In *SI Appendix* Fig. S9*A* we show that simulations of the individual-based model converge to exactly the same equilibria as in the main manuscript for $\upsilon = 0.1215$ and $\varphi = 0.15$. Finally, Fig. S9B shows the normalized effect of triploid fertility on the introgression of alleles through polyploid bridges. We see that normalizing for gamete frequency results in a faster introgression of alleles at low triploid fertility due to the increase in v and resulting higher frequency of tetraploids compared to when this is not normalized. Still, lower triploid fertility slows down introgression even without its effect on gamete equilibrium.

#### Text S3. Deterministic single-locus model

Let us consider a large panmictic population, where each organism bears a single biallelic locus with alleles $A$ and $a$, inhabiting a single deme with three cytotypes allowed to occur, i.e., diploids, triploids and tetraploids. Upon initialization the population has only diploids, whose genotype frequencies are given by the genotype vector $\boldsymbol{\gamma}_i^2(t)$, with $i \in \{AA, Aa, aa\}$ at generation $t$. The frequency of gamete types, both reduced and unreduced, formed following meiosis in a diploid organism are given in a gamete frequency matrix $G_2^{3\times5}$ of dimensions $3 \times 5$, where each row represents a genotype and each column represents a possible gamete, as follows:

$$
\begin{array}{cccccc}
 & A & a & AA & Aa & aa \\
\end{array}
$$

$$
G_2^{3\times5} = \begin{bmatrix} 1.0 & 0.0 & 1.0 & 0.0 & 0.0 \\ 0.5 & 0.5 & 0.25 & 0.5 & 0.25 \\ 0.0 & 1.0 & 0.0 & 0.0 & 1.0 \end{bmatrix} \begin{array}{c} AA \\ Aa \\ aa \end{array}
$$

Similarly, we define triploid and tetraploid genotype vectors to be $\boldsymbol{\gamma}_j^3(t)$ and $\boldsymbol{\gamma}_k^4(t)$, respectively, where $j \in \{AAA, AAa, Aaa, aaa\}$ and $k \in \{AAAA, AAAa, AAaa, aaa\}$. Their corresponding gamete frequency matrices are then:

$$
\begin{array}{cccccc}
 & A & a & AA & Aa & aa \\
\end{array}
$$

$$
G_3^{4\times5} \begin{bmatrix} 1.0 & 0.0 & 1.0 & 0.0 & 0.0 \\ 0.67 & 0.33 & 0.5 & 0.5 & 0.0 \\ 0.33 & 0.67 & 0.0 & 0.5 & 0.5 \\ 0.0 & 1.0 & 0.0 & 0.0 & 1.0 \end{bmatrix} \begin{array}{c} AAA \\ AAa \\ Aaa \\ aaa \end{array}
$$

And

$$
\begin{array}{cccccc}
 & A & a & AA & Aa & aa \\
\end{array}
$$

$$
G_4^{5\times5} \begin{bmatrix} 0.0 & 0.0 & 1.0 & 0.0 & 0.0 \\ 0.0 & 0.0 & 0.5 & 0.5 & 0.0 \\ 0.0 & 0.0 & 0.17 & 0.66 & 0.17 \\ 0.0 & 0.0 & 0.0 & 0.5 & 0.5 \\ 0.0 & 0.0 & 0.0 & 0.0 & 1.0 \end{bmatrix} \begin{array}{c} AAAA \\ AAAa \\ AAaa \\ Aaaa \\ aaaa \end{array}
$$

Notice that tetraploids are not allowed to produce haploid gametes, as there is no known mechanism of haploid gamete formation in tetraploid cytotypes. We then define the gamete pool of the population to be $\boldsymbol{\varpi} = \omega^2 + \omega^3 + \omega^4$, with $\omega^p = \gamma^p G_p, \forall\, p \in \{2,3,4\}$. The gamete pool from diploids is updated according to the probability of unreduced gametes $\upsilon = 0.10$ as follows: $\boldsymbol{\omega}_1 = (1-\upsilon)\boldsymbol{\omega}_1$, $\boldsymbol{\omega}_2 = (1-\upsilon)\boldsymbol{\omega}_2$, $\boldsymbol{\omega}_3 = (\upsilon)\boldsymbol{\omega}_3$, $\boldsymbol{\omega}_4 = (\upsilon)\boldsymbol{\omega}_4$ and $\boldsymbol{\omega}_5 = (\upsilon)\boldsymbol{\omega}_5$, where $\boldsymbol{\omega}_i$ is the *ith* element of $\varpi$. Then, we compute the zygote matrix $\zeta = \boldsymbol{\varpi}\boldsymbol{\varpi}^T$, where $T$ is the transpose of the vector $\boldsymbol{\varpi}$. As we do not distinguish between $Aa$ and $aA$, $\zeta$ is a symmetric matrix where each element $\zeta_{ij}$ represents the offspring's genotype generated for generation $t+1$. The reader shall easily verify that the new genotype vector for diploids $\boldsymbol{\gamma}_i^2(t+1)$ can be updated according to the following rule:

$$\boldsymbol{\gamma}_{AA}^2(t+1) = \zeta_{11}$$
$$\boldsymbol{\gamma}_{Aa}^2(t+1) = \zeta_{12} + \zeta_{21}$$
$$\boldsymbol{\gamma}_{aa}^2(t+1) = \zeta_{22}$$

The triploid genotype vector $\boldsymbol{\gamma}_j^3(t+1)$ then receives the information from the zygote matrix $\zeta$ decreased by a factor $\varphi = 0.30$, corresponding to triploid fertility (see Main Manuscript), as follows:

$$\boldsymbol{\gamma}_{AAA}^3(t+1) = \varphi(\zeta_{13} + \zeta_{31})$$
$$\boldsymbol{\gamma}_{AAa}^3(t+1) = \varphi(\zeta_{14} + \zeta_{23} + \zeta_{32} + \zeta_{41})$$
$$\boldsymbol{\gamma}_{Aaa}^3(t+1) = \varphi(\zeta_{15} + \zeta_{24} + \zeta_{42} + \zeta_{51})$$
$$\boldsymbol{\gamma}_{aaa}^3(t+1) = \varphi(\zeta_{25} + \zeta_{52})$$

and correspondingly the tetraploid genotype vector:

$$\boldsymbol{\gamma}_{AAAA}^4(t+1) = \zeta_{33}$$
$$\boldsymbol{\gamma}_{AAAa}^4(t+1) = (\zeta_{34} + \zeta_{43})$$
$$\boldsymbol{\gamma}_{AAaa}^4(t+1) = (\zeta_{35} + \zeta_{44} + \zeta_{53})$$
$$\boldsymbol{\gamma}_{Aaaa}^4(t+1) = (\zeta_{54} + \zeta_{45})$$
$$\boldsymbol{\gamma}_{aaaa}^4(t+1) = \zeta_{55}$$

The simulation consists of updating genotype frequencies iteratively for $T = 500$ generations. Notice that all genotype frequencies must be normalized at each generation so that the sum of genotype frequencies remains at 1 throughout the simulation. In *SI Appendix* Fig. S10*A* we can verify that the system behaves according to the assumptions of Hardy-Weinberg.

Finally, we assume two demes connected by migration at the tetraploid level controlled by a migration parameter $\kappa = 0.1$. We initialize one deme with one homozygous locus fixed for allele $A$ and a second deme with one homozygous locus fixed for allele $a$. Migration proceeds by taking a fraction $\kappa$ of tetraploid genotypes in one deme, and inserting it in the next deme, bidirectionally. Then, we can verify that the system converges to a stable equilibrium where each allele converges to a frequency of 50% in each deme (Fig. S10B).

##### Text S4. Derivation of the equations for the mainland-island model

We wish to study the speed with which a beneficial allele flowing from a mainland gets fixed in an island diploid gene pool. We first consider gene flow from tetraploids in the mainland into the island gene pool, which consists of diploids, triploids and tetraploids. Following the notation of the main manuscript, unidirectional gene flow occurs from the tetraploids in the mainland into the island with rate κ. Let us consider two alleles, $A$ and $a$, with frequencies in the island tetraploid gene pool $p_t$ and $q_t = 1 - p_t$ at time $t$, respectively. Then, five possible genotypes exist among the tetraploid cytotypes in the island; $AAAA$, $AAAa$, $AAaa$, $Aaaa$ and $aaaa$. At equilibrium, the frequency of each genotype is computed simply by the expansion of the binomial $(p+q)^4$, resulting in ${p_t}^4$ for $AAAA$, $4{p_t}^3 q_t$ for $AAAa$, $6{p_t}^2{q_t}^2$ for $AAaa$, $4p_t{q_t}^3$ for $Aaaa$ and ${q_t}^4$ for the homozygous $aaaa$. Assuming no double-reduction during meiosis, we can distinguish among three types of gametes $D_1 \triangleq AA$, $D_2 \triangleq Aa$ and $D_3 \triangleq aa$, produced with the following proportions in each genotype:

| **Genotype** | $\mathbf{D_1}$ | $\mathbf{D_2}$ | $\mathbf{D_3}$ |
|---|---|---|---|
| $AAAA$ | 1 | 0 | 0 |
| $AAAa$ | 1/2 | 1/2 | 0 |
| $AAaa$ | 1/6 | 2/3 | 1/6 |
| $Aaaa$ | 0 | 1/2 | 1/2 |
| $aaaa$ | 0 | 0 | 1 |

Now, we wish to compute the gamete types within the island produced by triploid genotypes. Let $r_t$ and $w_t$ denote the frequencies of allele $A$ and $a$, respectively, among the triploid genotypes. Then, assuming no double reduction, we identify five different gamete types, that is, two haploid gametes $H_1 \triangleq A$, $H_2 \triangleq a$ and three diploid gametes $D_1 \triangleq AA$, $D_2 \triangleq Aa$ and $D_3 \triangleq aa$, produced with the following proportions in each genotype:

| **Genotype** | $\mathbf{H_1}$ | $\mathbf{H_2}$ | $\mathbf{D_1}$ | $\mathbf{D_2}$ | $\mathbf{D_3}$ |
|---|---|---|---|---|---|
| $AAA$ | 1 | 0 | 1 | 0 | 0 |
| $AAa$ | 2/3 | 1/3 | 1/3 | 2/3 | 0 |
| $Aaa$ | 1/3 | 2/3 | 0 | 2/3 | 1/3 |
| $aaa$ | 0 | 1 | 0 | 0 | 1 |

As for the case of tetraploids, the frequency of each genotype is computed simply by the expansion of the binomial $(r_t + w_t)^3$, resulting in $r_t^{\,3}$ for $AAA$, $3r_t^{\,2}w_t$ for $AAa$, $3r_tw_t^{\,2}$ for $Aaa$ and $w_t^{\,3}$ for $aaa$. We assume triploids display reduced fertility $\varphi$ and produce haploid and diploid gametes with equal frequency. Then, we can update the frequency of diploid gametes in the population, following migration from the mainland and random mixing of diploid gametes produced by tetraploid and triploid genotypes as follows:

$$\mathrm{D}_1 = \{\mathrm{p_t}^4 + 2\mathrm{p_t}^3\mathrm{q_t} + \mathrm{p_t}^2\mathrm{q_t}^2\}\left(1 - \kappa - \frac{\varphi}{2}\right) + \kappa + \frac{1}{2}\varphi(\mathrm{r_t}^3 + \mathrm{r_t}^2\mathrm{w_t}) \qquad \text{S5.1}$$

$$\mathrm{D}_2 = \{2\mathrm{p_t}^3\mathrm{q_t} + 4\mathrm{p_t}^2\mathrm{q_t}^2 + 2\mathrm{p_t}\mathrm{q_t}^3\}\left(1 - \kappa - \frac{\varphi}{2}\right) + \frac{1}{2}\varphi(2\mathrm{r_t}^2\mathrm{w_t} + 2\mathrm{r_t}\mathrm{w_t}^2) \qquad \text{S5.2}$$

$$\mathrm{D}_3 = \{\mathrm{p_t}^2\mathrm{q_t}^2 + 2\mathrm{p_t}\mathrm{q_t}^3 + \mathrm{q_t}^4\}\left(1 - \kappa - \frac{\varphi}{2}\right) + \frac{1}{2}\varphi(\mathrm{r_t}\mathrm{w_t}^2 + \mathrm{w_t}^3) \qquad \text{S5.3}$$

That is, part of the diploid gametes produced within the island comes from triploid genotypes ($\varphi/2$), a frequency κ of gametes $D_1$ comes from the mainland tetraploid genotypes ($AAAA$), and the remaining is a result of gamete production by tetraploid genotypes within the island gamete pool. Moreover, notice that we simplify the system by assuming tetraploid genotypes are part of an established tetraploid population and do not depend on unreduced gamete frequencies from diploids.

We proceed in the same manner to recover the haploid gamete types within the island. Let $g_t$ and $z_t$ denote the frequency of the beneficial $A$ and the deleterious $a$ variants, respectively, and $H_1 \triangleq A$ and $H_2 \triangleq a$ the haploid gamete types within the island. A part $\varphi/2$ of haploid gametes come from the triploid genotypes within the island, and a part $\beta$ comes from the mainland diploid genotypes. Also, at equilibrium, each genotype is expected to occur according to the expansion of the binomial $(g_t + z_t)^2$, resulting in $\mathrm{g_t}^2$ for the genotype $AA$, $2\mathrm{g_t}\mathrm{z_t}$ for the genotype $Aa$ and $\mathrm{z_t}^2$ for $aa$. Again, assuming no double-reduction, we write:

$$\mathrm{H}_1 = \{\mathrm{g_t}^2 + \mathrm{g_t}\mathrm{z_t}\}\left(1 - \beta - \frac{\varphi}{2}\right) + \beta + \frac{1}{2}\varphi(\mathrm{r_t}^3 + 2\mathrm{r_t}^2\mathrm{w_t} + \mathrm{r_t}\mathrm{w_t}^2) \qquad \text{S5.4}$$

$$\mathrm{H}_2 = \{\mathrm{g_t}\mathrm{z_t} + \mathrm{z_t}^2\}\left(1 - \beta - \frac{\varphi}{2}\right) + \frac{1}{2}\varphi(\mathrm{r_t}^2\mathrm{w_t} + 2\mathrm{r_t}\mathrm{w_t}^2 + \mathrm{w_t}^3) \qquad \text{S5.5}$$

The frequency of each genotype in each cytotype within the island is then obtained by the expansion $(D_1 + D_2 + D_3 + \mathrm{H}_1 + \mathrm{H}_2)^2$ followed by additive selection $s$ on the deleterious mutations:

Tetraploid – $AAAA$: $\frac{1}{\Gamma} {D_1}^2 (1 - 0s)$

Tetraploid – $AAAa$: $\frac{1}{\Gamma} 2D_1D_2 \left(1 - \frac{1}{4}s\right)$

Tetraploid – $AAaa$: $\frac{1}{\Gamma} \left\{{D_2}^2 + 2D_1D_3\right\} \left(1 - \frac{1}{2}s\right)$

Tetraploid – $Aaaa$: $\frac{1}{\Gamma} 2D_2D_3 \left(1 - \frac{3}{4}s\right)$

Tetraploid – $aaaa$: $\frac{1}{\Gamma} {D_3}^2 (1 - s)$

Triploid – $AAA$: $\frac{1}{\Gamma} 2H_1D_1 (1 - 0s)$

Triploid – $AAa$: $\frac{1}{\Gamma} \{2\mathrm{H}_1\mathrm{D}_2 + 2\mathrm{H}_2\mathrm{D}_1\} \left(1 - \frac{1}{3}s\right)$

Triploid – $Aaa$: $\frac{1}{\Gamma} \{2\mathrm{H}_1\mathrm{D}_3 + 2\mathrm{H}_2\mathrm{D}_2\} \left(1 - \frac{2}{3}s\right)$

Triploid – $aaa$: $\frac{1}{\Gamma} 2\mathrm{H}_2\mathrm{D}_3 (1 - s)$

Diploid – $AA$: $\frac{1}{\Gamma} {H_1}^2 (1 - 0s)$

Diploid – $Aa$: $\frac{1}{\Gamma} 2\mathrm{H}_1\mathrm{H}_2 \left(1 - \frac{1}{2}s\right)$

Diploid – $aa$: $\frac{1}{\Gamma} {H_2}^2 (1 - s)$

with,

$$\Gamma = {D_1}^2 + 2D_1D_2 \left(1 - \frac{1}{4}s\right) + \left\{{D_2}^2 + 2D_1D_3\right\} \left(1 - \frac{1}{2}s\right) + 2D_2D_3 \left(1 - \frac{3}{4}s\right) + {D_3}^2 (1 - s) + 2H_1D_1 + \{2\mathrm{H}_1\mathrm{D}_2 + 2\mathrm{H}_2\mathrm{D}_1\} \left(1 - \frac{1}{3}s\right) + \{2\mathrm{H}_1\mathrm{D}_3 + 2\mathrm{H}_2\mathrm{D}_2\} \left(1 - \frac{2}{3}s\right) + 2\mathrm{H}_2\mathrm{D}_3 (1 - s) + {H_1}^2 + 2\mathrm{H}_1\mathrm{H}_2 \left(1 - \frac{1}{2}s\right) + {H_2}^2 (1 - s)$$

Now, we can write the recursive equations for the frequency of $A$ in time for each cytotype within the island, by recovering the frequency of the beneficial mutation in each of the genotypes outlined above. For the frequency of $A$ among the tetraploids we have:

$$p_{t+1} = \frac{1}{\Lambda} \left\{ {D_1}^2 + \frac{3}{2}{D_2}^2 + 2D_1D_3 \left(1 - \frac{1}{4}s\right) + \frac{1}{2}\left\{{D_2}^2 + 2D_1D_3\right\} \left(1 - \frac{1}{2}s\right) + \frac{1}{2}D_2D_3 \left(1 - \frac{3}{4s}\right) \right\} \qquad \text{S5.6}$$

where,

$$\Lambda = {D_1}^2 + 2D_1D_2 \left(1 - \frac{1}{4}s\right) + \left\{{D_2}^2 + 2D_1D_3\right\} \left(1 - \frac{1}{2}s\right) + 2D_2D_3 \left(1 - \frac{3}{4}s\right) + {D_3}^2 (1 - s)$$

The frequency of $A$ among triploid and diploid genotypes are computed as follows:

$$r_{t+1} = \frac{1}{\Phi}\left\{2H_1D_1 + \frac{2}{3}\{2\mathrm{H}_1\mathrm{D}_2 + 2\mathrm{H}_2\mathrm{D}_1\}\left(1 - \frac{1}{3}s\right) + \frac{1}{3}\{2\mathrm{H}_1\mathrm{D}_3 + 2\mathrm{H}_2\mathrm{D}_2\}\left(1 - \frac{2}{3}s\right)\right\} \quad \text{S5.7}$$

where,

$$\Phi = 2H_1D_1 + \{2\mathrm{H}_1\mathrm{D}_2 + 2\mathrm{H}_2\mathrm{D}_1\}\left(1 - \frac{1}{3}s\right) + \{2\mathrm{H}_1\mathrm{D}_3 + 2\mathrm{H}_2\mathrm{D}_2\}\left(1 - \frac{2}{3}s\right) + 2\mathrm{H}_2\mathrm{D}_3(1 - s)$$

and,

$$g_{t+1} = \frac{1}{\Psi}\left\{{H_1}^2 + H_1H_2\left(1 - \frac{1}{2}s\right)\right\} \quad \text{S5.8}$$

with,

$$\Psi = {H_1}^2 + 2H_1H_2\left(1 - \frac{1}{2}s\right) + {H_2}^2(1 - s)$$

#### Text S5. Recursive nonlinear equations with triploid gametes

Here, we modify the set of equations $2.1 - 2.5$ in the main manuscript to include triploid gametes. The assumptions are the same as in the main manuscript, except that now, with the inclusion of triploid gametes, pentaploid and hexaploid genotypes could also be generated. As our knowledge of gamete production in these genotypes is too restricted, and they often display very low viability, we assume such genotypes to be lethal, and therefore they do not contribute to the gamete pool in our system. The new set of equations describing cytotype dynamics on a large population can then be written as follows:

$$g_1 = x_2(t)(1-\upsilon) + \lambda_1 x_3(t)\varphi \quad \text{S4.1}$$

$$g_2 = x_2(t)\upsilon + \lambda_2 x_3(t)\varphi + x_4(t) \quad \text{S4.2}$$

$$g_3 = \lambda_3 x_3(t)\varphi \quad \text{S4.3}$$

where $G_3$, now, corresponds to the proportion of triploid gametes produced in the system, and $\lambda_1$, $\lambda_2$ and $\lambda_3$ the frequency of haploid, diploid and triploid gametes produced by triploid genotypes. Then, the genotype frequencies can be updated as follows:

$$\mathrm{x}_2(\mathrm{t}+1) = \frac{{g_1}^2}{\Omega} \quad \text{S4.4}$$

$$\mathrm{x}_3(\mathrm{t}+1) = \frac{2g_1g_2}{\Omega} \quad \text{S4.5}$$

$$\mathrm{x}_4(\mathrm{t}+1) = \frac{{g_2}^2 + 2g_1g_3}{\Omega} \quad \text{S4.6}$$

with $\Omega = (g_1 + g_2 + g_3)^2$ as before.

From this set of equations, it becomes clear that the only change concerns the higher load of tetraploid genotypes in the population for every time t, given by the new term $2g_1g_3$ in S1.6. For $\lambda_1 = \lambda_2 = 0.25$ and $\lambda_3 = \lambda_1 + \lambda_2$, we verify the same pattern of cytotype dynamics as given by the set of Eq. $4.1 - 4.5$ in the main manuscript, except that polyploids overtake the system with considerably lower values of $\bar{\upsilon}$, as expected (*SI Appendix* Fig. S11*A*).

To have a clearer understanding of why self-fertilization of triploids leads ultimately to tetraploid genotypes let us consider a triploid plant that produces a large number of pollen. As Cao et al. (1) do not provide the frequency of pollen ploidy levels produced by their systems, let us assume that gametes follow the average frequencies computed by Ramsey and Schemske (2). Then, a triploid plant shall produce $G_1 = 3\%$, $G_2 = 2\%$ and $G_3 = 5.2\%$ of haploid, diploid and triploid gametes, respectively, with all remaining gametes being aneuploid, which we assume to be not viable for simplicity. If self-fertilization occurs randomly, then we can compute the frequency of genotypes among the offspring in the F2 generation as:

$$hexaploids = \frac{{G_3}^2}{\Omega} = 0.2599$$

$$pentaploids = \frac{2\, G_3 G_2}{\Omega} = 0.1999$$

$$tetraploids = \frac{2\, G_3 G_1 + {G_2}^2}{\Omega} = 0.3383$$

$$triploids = \frac{2 G_1 G_2}{\Omega} = 0.1153$$

$$diploids = \frac{{G_1}^2}{\Omega} = 0.0865$$

with $\Omega = (G_1 + G_2 + G_3)^2$. Clearly, the frequency of polyploids (3x or higher) is much more prevalent in only one generation. Unfortunately, our knowledge of gamete ploidy levels produced by pentaploids and hexaploids is too restricted, which prevents us from calculating safely the genotype frequencies of succeeding generations. However, diploid gametes tend to be much more frequent in higher ploidy cytotypes, and are the standard ploidy level in gametes from tetraploids (3). Thus, it is easy to see that tetraploid frequency can only increase over succeeding generations, especially if pentaploids and hexaploids are highly unstable genotypes. The latter scenario can also be seen by iterating equations $S2.1 - S2.6$ with initial conditions $x_2(0) = 0, x_3(0) = 1$ and $x_4(0) = 0$ (Fig. S11*B*). Moreover, in one of the strains used by the authors, only 4 plants were selected from the F2 (hybrid triploids) progeny to produce the F3 progeny. Based our simple calculations, the expected number of diploid organisms found would be simply $0.0865(4) = 0.346$.

#### Algorithm S1. Implementation of the mixed-ploidy stochastic model of interspecific gene flow

Here, we describe the technical details for implementation of the main model used in the manuscript. This model is a special case of the deterministic single locus model presented in Text S3. The main difference being drift due to sampling of finite population sizes and many loci. This work has been carried out in Java, and below we provide a detailed explanation of the code rationale, which can be easily implemented in alternative programming languages. The project consists of three classes: *Bridge*, *Evolution* and *StartGenome*. The class *Bridge* implements the main structure of the model, where an initial population is called, and generations are built iteratively, with bidirectional migration at the end of each generation. The class *Evolution* implements the algorithm for meiosis described within the main manuscript, and additional helping methods for measuring allele frequencies. Finally, *StartGenome* initializes an initial population where the genome configuration of organisms are instantiated. Let us start by the class *Bridge*.

```
//We start with the parameters that control our simulation.
int generations = 20000; //Number of generations for which the simulation must run
int popSize = 1000; //Carrying capacity of the system
int chromLength = 100; //Length of the chromosome, i.e., number of unlinked loci
double meiosisCoeff = 0.05; //Proportion of loci that undergo recombination during meiosis
double probPolyploid = 0.1; //Probability that meiosis in diploids will produce diploid gametes
double reducedFertilityTriploid = 0.70; //One minus triploid fertiliy
double kappa = 0.1; //Migration rate at the tetraploid level

/*Then we create two instances of the classes StartGenome and Evolution.
These instances will allow us to initialize a population, and call the appropriate methods.*/

StartGenome initiate = new StartGenome(popSize, chromLength);
Evolution resolve = new Evolution(meiosisCoeff, probPolyploid, reducedFertilityTriploid);

/*Initial populations are stored in two HashMaps, so we can easily retrieve the genome of individuals by
their indexes, given by Integers. Pool01 will refer to deme A, whereas pool02 refers to deme B. startPop is
a method inside class StartGenome that receives as parameters the deme number, and generates a
population of size popSize of diploid individuals whose chromosomes have chromLength loci. We will
provide details about the class later.*/

HashMap<Integer, int[][]> pool01 = initiate.startPop(1);
HashMap<Integer, int[][]> pool02 = initiate.startPop(2);

/*Now, we create a while loop for the number of generations. For simplicity, we describe the procedure
for only one deme, which is exactly the same for the case of two demes. */

int t = 0;
while(t < generations) {

    //Initialize two empty HashMaps that will store the next generation
```

```
        HashMap<Integer, int[][]> newPool01 = new HashMap<>();

        int offspringNumber = 0; //We control the offspring number to fill the HashMaps in order.
        //While the new population is below carrying capacity
        while(newPool01.size() < popSize) {

        //Choose two random mates from the original population
        int mate01 = (int) (Math.random()*pool01.size());
        int mate02 = (int) (Math.random()*pool01.size());

        //We need to control for a possible null value, owing to triploid fertility.
            try{
            //Call method meiosis in class Evolution for both parents
            int[][] gamete01 = resolve.meiosis(pool01.get(mate01));
            int[][] gamete02 = resolve.meiosis(pool01.get(mate02));
            //Then join gametes to produce offspring
            int[][] offspring = joinGametes(gamete01, gamete02);
            //Store offspring for the next generation
            newPool01.put(offspringNumber, offspring);
                    offspringNumber ++;
                    }catch(NullPointerException e) {

                    }
            }
            pool01 = newPool01; //Update original population
        }
/*With two demes, random tetraploid individuals are taken from pool01 and placed into pool02, and vice versa. Each
tetraploid has probability kappa of migrating.*/
t = t+1;
}
```

Now, we described the Meiosis method in the class *Evolution*. We use switch statements for the cases of diploid (2), triploid (3) and tetraploid (4) genotypes. All cases operate in the same way within the software, except for the different chromosome numbers being handled. Here, we describe how the method handles diploid genotypes, which can be easily extended to triploid and tetraploid genotypes, see main manuscript and *SI Appendix* Fig. S4.

```
/*The method receives the individual's genome given in a two dimensional vector with n rows (ploidy) and L columns
(loci). It recombines numAlleles random loci and with probability probPolyploid created unreduced gametes*/

public int[][] meiosis(int[][] genome) {
                    //We compute the number of alleles that must recombine.
            int numAlleles = (int) (genome[0].length*this.meiosisCoeff);
            int[][] gametes = null; //initialize an empty gamete.

            switch (genome.length) {
            case 2: //Diploidy
                    // Exchanges alleles on random loci. The number of loci is given by meiosisCoeff*genome.length.

                    for(int i = 0; i < numAlleles; i++) {
                                    // Find a random locus and exchange alleles
```

```
                int locus = (int) (Math.random()*genome[0].length);
                int key = genome[0][locus];
                genome[0][locus] = genome[1][locus];
                genome[1][locus] = key;
            }
            double r = Math.random();
    //probPolyploid is the frequency of unreduced gametes given in the constructor of the Class.
            if(r < this.probPolyploid) {
                //initialize a diploid gamete and copy chromosomes
                gametes = new int[2][genome[0].length];
                for(int i = 0; i < genome[0].length; i++) {
                    gametes[0][i] = genome[0][i];
                    gametes[1][i] = genome[1][i];
                }
            }else {
                //If not unreduced, initialize a one-dimensional vector
                gametes = new int[1][genome[0].length];
                double pair = Math.random();
                //Select one of the chromosomes with equal probability
                if(pair < 0.5) {
                    for(int i = 0; i < genome[0].length; i++) {
                        gametes[0][i] = genome[0][i];
                    }
                }else {
                    for(int i = 0; i < genome[0].length; i++) {
                        gametes[0][i] = genome[1][i];
                    }
                }
            }
            break;
        }
//Then gametes are returned.
return gametes;
}
```

Now, the class *StartGenome* initializes the population. Here, we describe for the case when each pool is fixed for one different alleles at every loci.

```
public class StartGenome {

    private int popSize; //Population size of each deme
    private int chromLength; //Number of loci in each organism
    private HashMap<Integer, int[][]> population; //Stores the population

//We have a public constructor that receives the 2 parameters
    public StartGenome(int popSize, int chromLength) {

        this.popSize = popSize;
        this.chromLength = chromLength;
    }

    public HashMap<Integer, int[][]> startPop(int deme) {
```

```
		HashMap<Integer, int[][]> population = new HashMap<>();

// Case 1 corresponds to deme A and case 2 corresponds to deme B
		switch(deme) {
		case 1:
//We initialize a diploid genome with chromLength loci fixed with allele 1
				for(int i = 0; i < this.popSize; i++) {
					int[][] genome = new int[2][this.chromLength];
					for(int j = 0; j < this.chromLength; j++) {
						genome[0][j] = 1;
						genome[1][j] = 1;
					}
//Every genome is stored in the HashMap which is manipulated within the class Bridge
					population.put(i, genome);
				}
				break;
		case 2:
				for(int i = 0; i < this.popSize; i++) {
					int[][] genome = new int[2][this.chromLength];
					for(int j = 0; j < this.chromLength; j++) {
						genome[0][j] = 0;
						genome[1][j] = 0;
					}
					population.put(i, genome);
				}
				break;
		}
//Finally, we return the population.
		return population;
	}

}
```

Finally, the class *StartGenome* has a default constructor that takes as parameters the population size and chromosome length, as follows:

```
public StartGenome(int popSize, int chromLength) {

		this.popSize = popSize;
		this.chromLength = chromLength;
	}
```

Then, an initial population is constructed using the public method startPop():

```
	public HashMap<Integer, int[][]> startPop(int island) {

		/*We initialize the population using a HashMap, where the first element is the individual index and the two-
dimensional vector "int[][]" stores the genome of the individual. Each element of the rows correspond to a
chromosome, whereas elements of the columns correspond to loci. So, for example, int[1][2] identifies the second
locus of the 1st chromosome. */

		HashMap<Integer, int[][]> population = new HashMap<>();
```

```
    //We use switch statements for the population of each deme (island)
    // Island = 1 receives genomes fixed for allele 1, whereas Island = 2 receives genomes fixed for allele 0.
    switch(island) {
    case 1:
            for(int i = 0; i < this.popSize; i++) {
                    int[][] genome = new int[2][this.chromLength]; // diploid - length 2
                    for(int j = 0; j < this.chromLength; j++) {
                            genome[0][j] = 1;
                            genome[1][j] = 1;
                    }
                    population.put(i, genome);
            }
            break;
    case 2:
            for(int i = 0; i < this.popSize; i++) {
                    int[][] genome = new int[2][this.chromLength];
                    for(int j = 0; j < this.chromLength; j++) {
                            genome[0][j] = 0;
                            genome[1][j] = 0;
                    }
                    population.put(i, genome);
            }
            break;
    }
    return population;
}
```

#### Algorithm S2. Retrieval of $\kappa_{min} \leftarrow (s, \beta)$

For simplicity, we will refer to Equations presented within this Supporting Information file, rather than to equations described within the main manuscript. The algorithm below receives as parameters $s$ and $\beta$. It is executed for all values of $s \in [0, 0.10]$ and $\beta \in (0, 0.05]$ to produce the results in the main manuscript.

```
/* Initialize allele frequencies within the island diploid pool */
g_t = 0.0, z_t = 1.0 − 0.0
/* Initialize time step */
t = 0
/* Initiliaze time for full introgression of the beneficial variant at
the diploid level within the island to occur */
κ_min ← 0 /* No gene flow at the tetraploid level */
φ ← 0 /* No triploid interference */
t_max = 0
while(t_max == 0) do
        /* Update gamete types within the island diploid gene pool */
        H_1 ← S4.4
        H_2 ← S4.5
        /* Update frequency of the beneficial variant */
        g_t ← S4.8
        if(g_t < 1.0) then
            t ← t + 1
        else
            t_max ← t
            break
        end
end
/* Set tetraploid − level introgression rate to β */
κ_min ← β, φ ← 0.30, β ← 0
while(κ_min ≤ 1.0) do
        t = 0
        g_t = 0.0, z_t = 1.0 − 0.0
        p_t = 0.0, q_t = 1.0 − 0.0
        r_t = 0.0, w_t = 1.0 − 0.0
        while(t < t_max) do
                /* Update gamete types within the island gene pool */
                D_1 ← S4.1
                D_2 ← S4.2
                D_3 ← S4.3
                H_1 ← S4.4
                H_2 ← S4.5
                g_t ← S4.8
                t ← t + 1
        end
        if(g_t < 1.0) then
            κ_min ← κ_min + 10^-6 /* Sets κ_min resolution */
        else
            break
        end
end
return κ_min
```

#### Supporting Figures

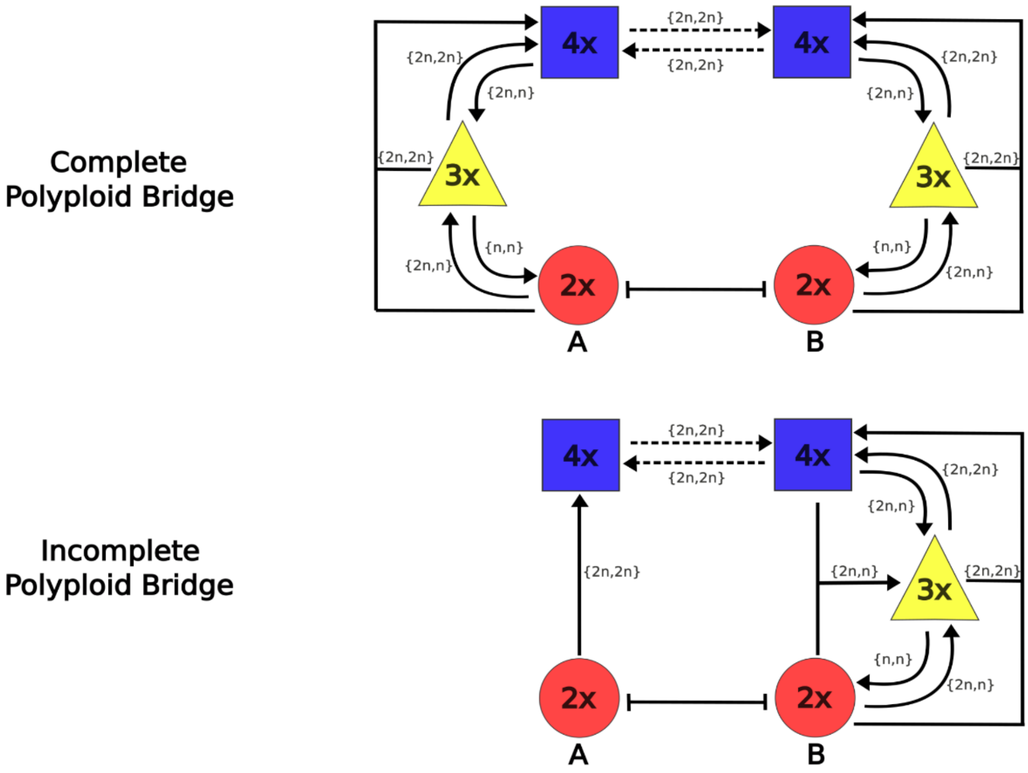

**Fig. S1.** Complete and incomplete polyploid bridges. Triploids might inhabit contact zones between diploid and tetraploid populations, and therefore be the source of gene transfer from tetraploids to back to diploids inside one mixed-ploidy population, or be completely absent in one of the species, leading to asymmetrical gene flow. A third possibility is hybridization between diploids in population A directly with tetraploids in population B. Such a phenomenon has been described in the literature, but is not within the scope of this study.

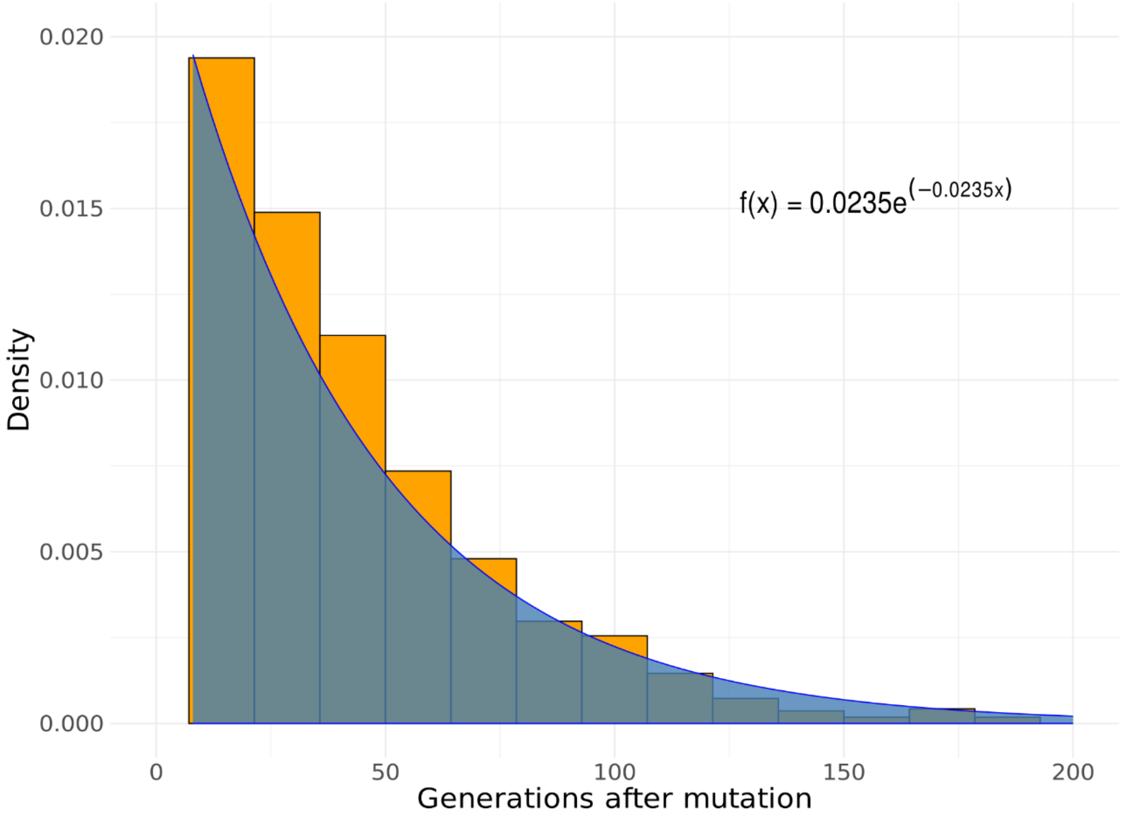

**Fig. S2.** Time to appearance of mutant allele in population B. We ran 10000 simulations and measured the time it takes for a mutant allele in the diploid level of population A to appear at the diploid level of population B. The distribution of waiting times is well described by an exponential distribution with an expected value of $\sim 43$ generations (see Equation in the inset). Parameters: $\upsilon = 0.10$, $\varphi = 0.30$, $\kappa = 0.30$.

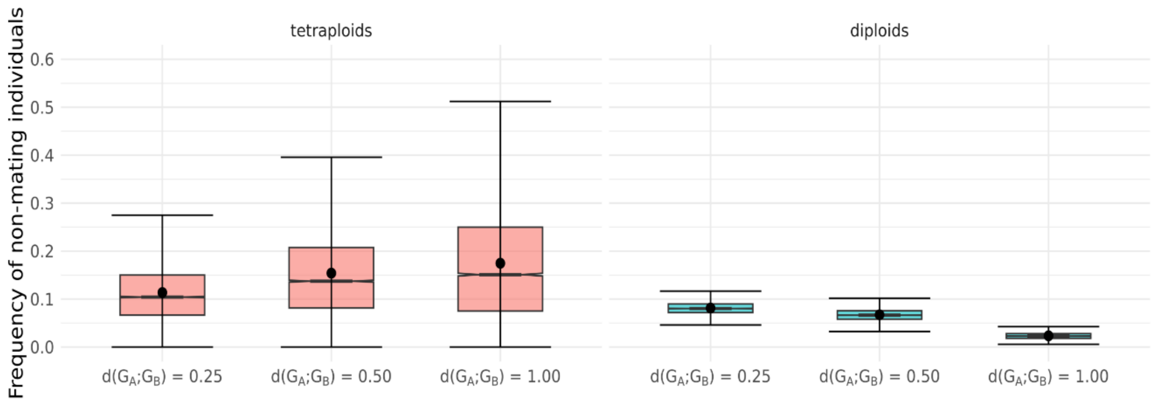

**Fig. S3**. Frequency of non-mating cytotypes for various genetic distances between demes A and B. Notice that as the genetic divergence between demes increases, the frequency of tetraploid cytotypes that do not mate successfully increases, and thus gene flow down to the diploid state cannot happen. This explains why the frequency of non-mating diploid cytotypes, on the other hand, decreases as genetic divergence increases, i.e., deleterious alleles cannot penetrate the diploid state when divergence is too high.

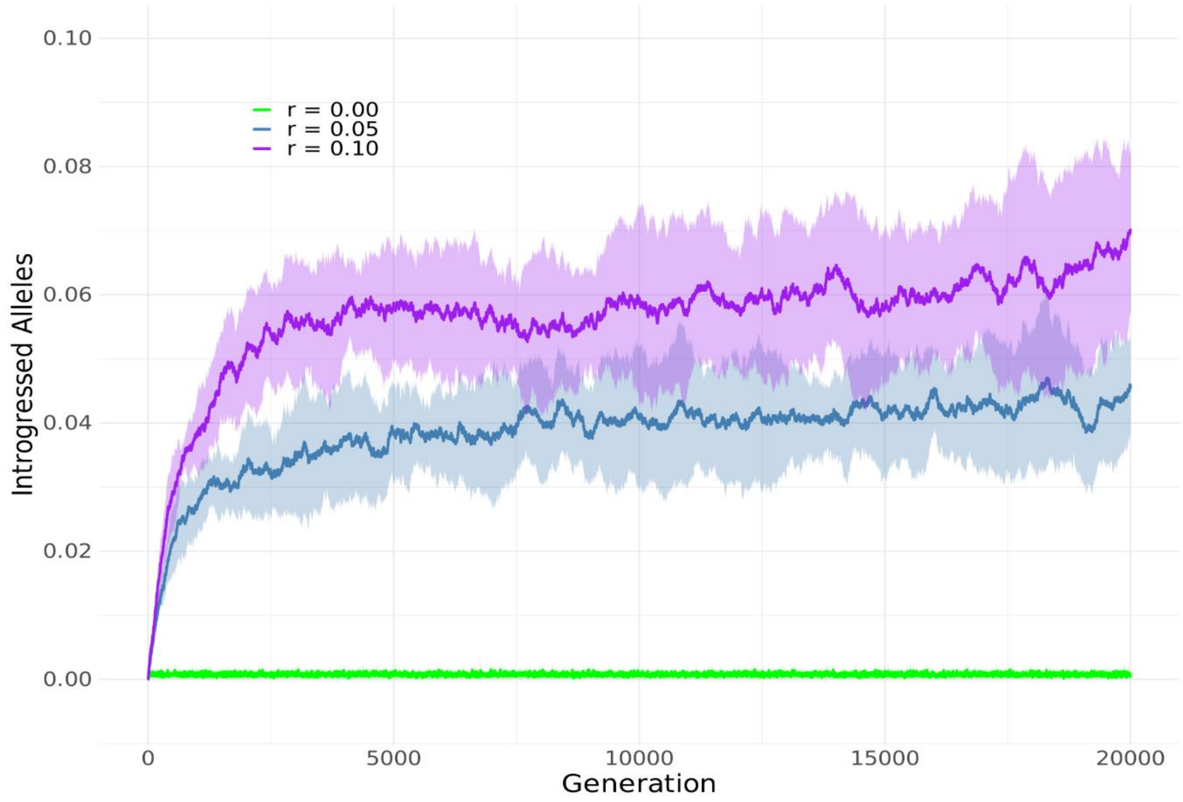

**Fig. S4.** Effect of recombination rate (r) on the average frequency of introgressed alleles in both mixed-ploidy populations.

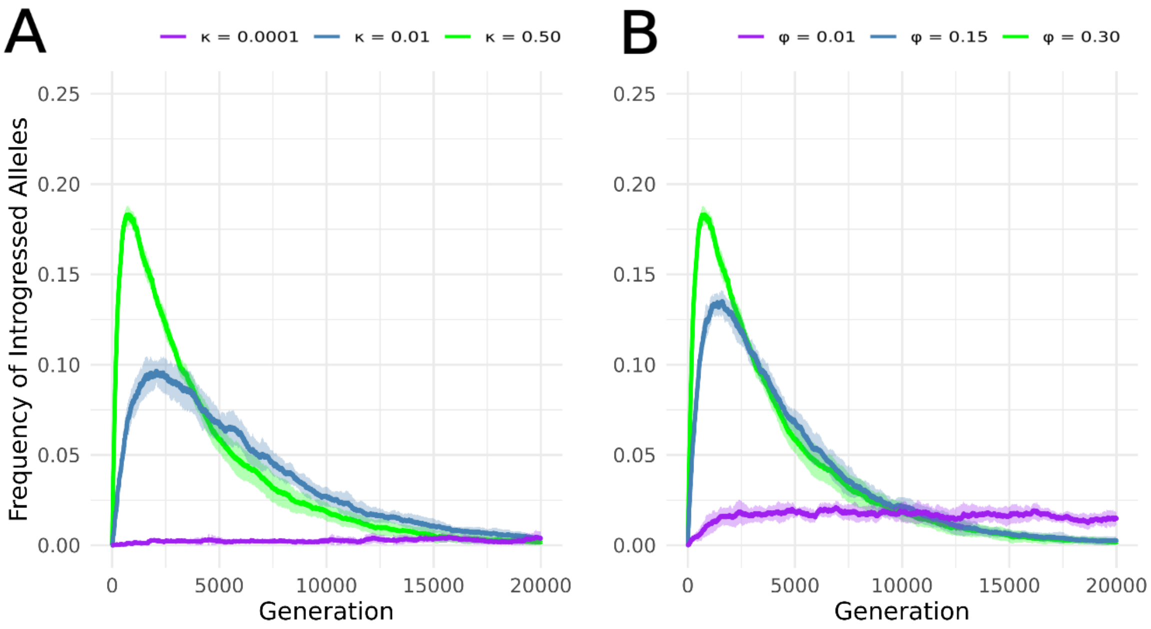

**Fig. S5**. Frequency of introgressed alleles in heterozygous loci. Frequency of introgressed alleles in loci where one of the alleles is native. Notice that introgression of non-native alleles proceeds by first sharing loci with native alleles and then decays rather quickly, because of full introgression, or fixation (see manuscript).

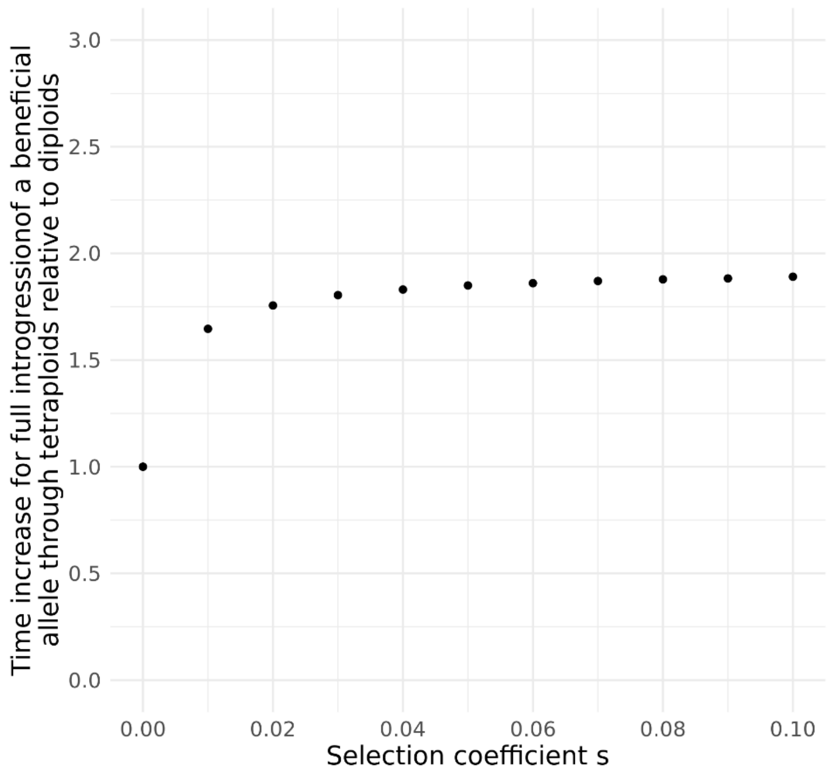

**Fig. S6.** Increase in time necessary for introgression through tetraploids to be as efficient as introgression through diploids. Ratio is shown of the time for fixation of the beneficial allele in the island tetraploid gene pool (through the route 4x → 4x) over the time for fixation of the beneficial allele in the island diploid gene pool (through the route 2x → 2x). For a neutral allele (s = 0) the speed of introgression does not differ between the two cases. As the allele coming from the mainland becomes increasingly beneficial (s > 0) the relative time for introgression between tetraploid gene pools compared to diploid gene pools approaches 2. In other words, with increasing s gene flow between tetraploids is twice less efficient compared to introgression between diploids.

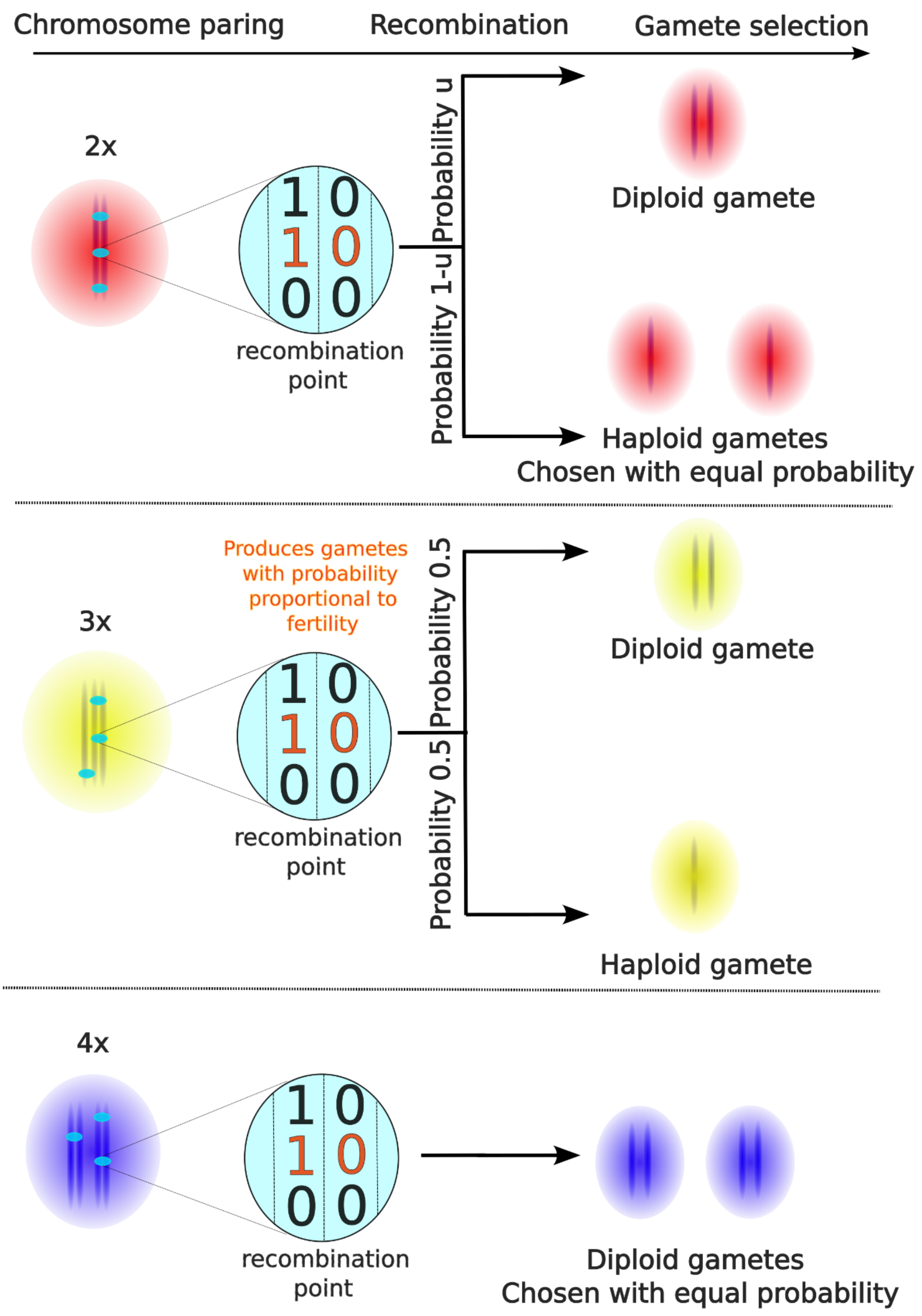

**Fig. S7.** Schematic representation of the computational procedure used to represent meiosis and gamete formation for diploids, triploids and tetraploids, in our model.

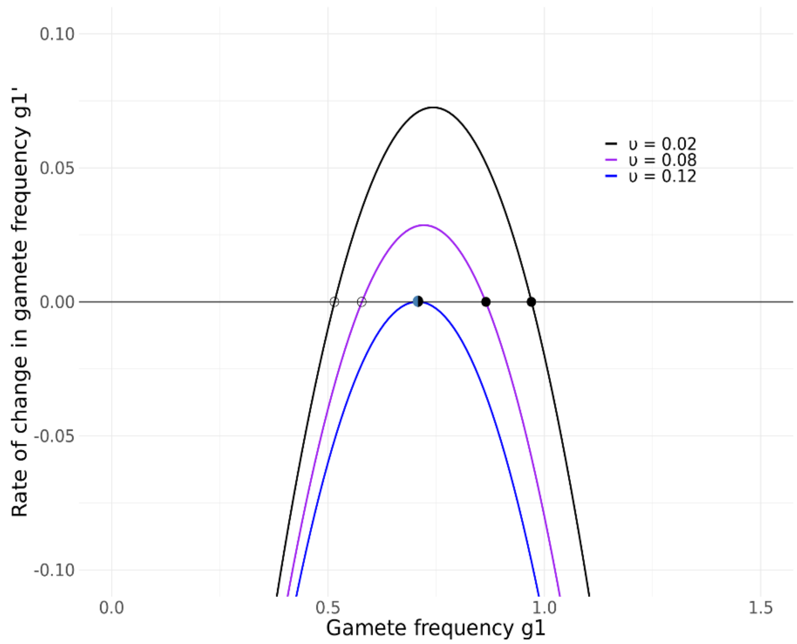

**Fig. S8. Phase portrait of the dynamical system derived from a mixed-ploidy panmictic unit.** The phase portrait is constructed by Eq. S1.5, with $\varphi = 0.30$, and three different values of unreduced gametes frequency $\upsilon$. Stable and unstable points are given by filled and open circles, respectively. The half-stable point (steel blue and black) at $\upsilon = 0.12$ represents the bifurcation point where gamete frequency $g_1$ converges to zero from its initial frequency of 100% following the initial conditions of the system. Bifurcation points can be calculated by S1.8 for any $\varphi$.

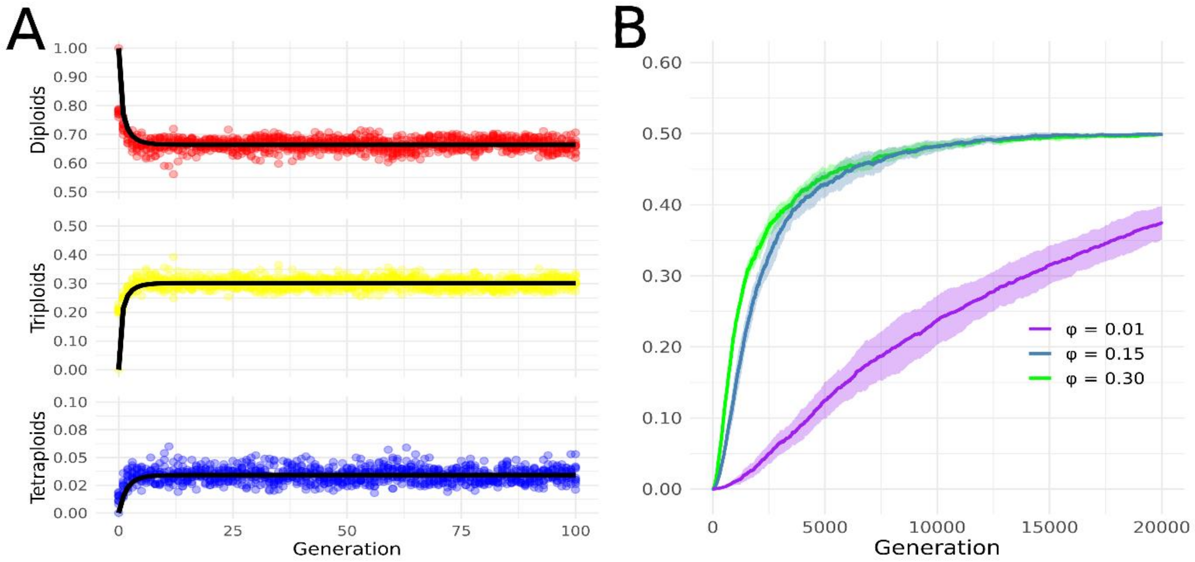

**Fig. S9.** Cytotype dynamics and effect of triploid fertility on introgression with normalized frequency of unreduced gametes**.** (*A*) Both the numerical simulations with the set of Eqs. $1-2$ in the main manuscript, with parameters $\varphi = 0.15$ and $\upsilon = 0.1215$. The equilibria for cytotype frequencies are the same as in the main manuscript. Dots correspond to the individual-based model with finite population size (S=1000), and black solid lines to the iterated maps. (*B*) Effect of fertility shown to be the same as in the main manuscript.

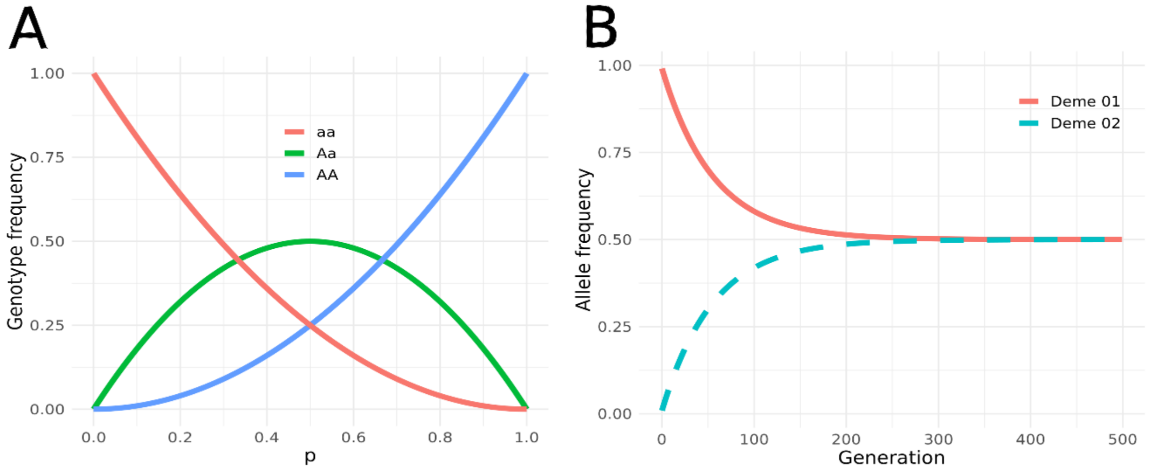

**Fig. S10.** WGD-mediated introgression dynamics in the deterministic single-locus model**.** (*A*) Convergence of our model to an equilibrium as expected from Hardy-Weinberg populations. The initial frequency of the allele A is denoted by p, and equilibrium is measured at the diploid level. (B) Unfolding of introgression in time given migration at the tetraploid level for $\kappa = 0.1$.

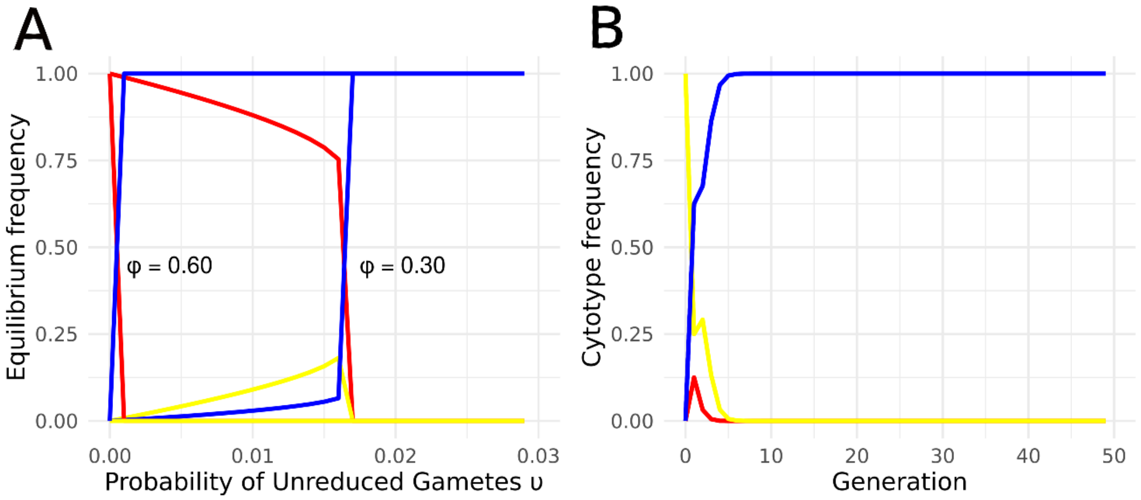

**Fig. S11.** Cytotype dynamics in a single deme with triploid gametes**.** (*A*) Equilibrium points given by Eqs. $S1.1 - S1.6$. Compared with equilibria in the main manuscript, we notice that the frequency of unreduced gametes required for tetraploids to overtake the system is considerably lower, for the same triploid fertilities. This is due the new source of gametes (3n) that can only produce higher ploidy taxa, thus supressing the frequency of diploids. (*B*) Numerical simulations of Eqs. $S1.1 - S1.6$ with only triploids in the start of the simulation. Clearly, tetraploids overtake the system quite rapidly, as diploids are rarely produced and rapidly replaced as discussed in *SI Appendix Text S2*.

### REFERENCES

1. Y. Cao *et al.*, Genome balance and dosage effect drive allopolyploid formation in Brassica. *Proc Natl Acad Sci U S A* **120**, e2217672120 (2023).
2. J. R. and, D. W. Schemske, PATHWAYS, MECHANISMS, AND RATES OF POLYPLOID FORMATION IN FLOWERING PLANTS. *Annual Review of Ecology and Systematics* **29**, 467-501 (1998).
3. B. Arnold, S. T. Kim, K. Bomblies, Single Geographic Origin of a Widespread Autotetraploid Arabidopsis arenosa Lineage Followed by Interploidy Admixture. *Mol Biol Evol* **32**, 1382-1395 (2015).
